# Transcription elongation factor SPT6L recruits ARGONAUTE to guide mRNA cytosine methylation preventing premature termination in plants

**DOI:** 10.64898/2026.03.25.714125

**Authors:** Tomáš Kašpar, Vojtěch Čermák, Kateřina Adamusová, Lukáš Fischer

## Abstract

In plants, an essential component of the RNA Polymerase II (Pol II) complex, SPT6L, contains a unique C-terminal ARGONAUTE binding domain - AGO-hook. Although it has been recently shown that SPT6L is also part of the Pol V complex, where its AGO-hook might participate in the RNA-directed DNA methylation pathway, several lines of evidence suggest that the AGO-hook should also have a function in the Pol II complex. Here we demonstrate that in *Arabidopsis thaliana*, SPT6L recruits AGO4 via the AGO-hook domain, independently of Pol V. Using direct RNA sequencing, we show that this interaction is connected with guided deposition of 5-methylcytosine (m^5^C) on Pol II transcripts. The role of AGO4 in m^5^C guidance was further supported by increased methylation of *GFP* transcripts after their targeting by sRNAs in tobacco BY-2 cells. Removing the SPT6L AGO-hook (*spt6lΔAh*) in *A. thaliana* resulted in the hypomethylation of specific m^5^C sites, mimicking even stronger RNA hypomethylation observed in the *ago4* mutant. Native elongating transcript sequencing revealed that impaired targeting of m^5^C methylation in the *spt6lΔAh* mutant was accompanied by Pol II stalling downstream of hypomethylated cytosines. Furthermore, this stalling appeared to be associated with premature transcription termination, which we also detected in published transcriptomes of mutants lacking AGO4, as well as TRM4B that is the major RNA m^5^C methyltransferase. Our findings uncover a novel “guide-and-modify” mechanism in which SPT6L integrates sRNA-directed epitranscriptomic marking with transcription elongation, ensuring proper mRNA termination.

## Introduction

In all eukaryotes, Suppressor of Ty insertion 6 (SPT6) proteins are a canonical part of the Pol II complex, functioning as histone chaperones and transcription elongation factors. During the evolution of plants, an ortholog SPT6-like (SPT6L) acquired a disordered domain enriched in WG/GW motifs at the C-terminal end (Karlowski et al., 2010; Čermák et al., 2024). Such domains were shown to function as an ARGONAUTE (AGO) binding platform - an AGO-hook (El-Shami et al., 2007; Karlowski et al., 2010; Zielezinski and Karlowski, 2015).

In plants, other AGO-hook-containing proteins have been shown to recruit AGO complexes with small RNA (sRNA) to guide DNA methylation in the process called RNA-directed DNA methylation (RdDM). These proteins are NUCLEAR RNA POLYMERASE E1 (NRPE1), the largest subunit of Pol V complex, and its transcription elongation factor SPT5L, which is a paralogue of the Pol II-associated SPT5 protein (El-Shami et al., 2007; Bies-Etheve et al., 2009; Trujillo et al., 2016). Besides the function of AGO-hooks in RdDM, there have been others with various functions: NEEDED FOR RDR2-INDEPENDENT DNA METHYLATION (NERD) protein binds protein AGO2 to promote chromatin-based silencing pathways in plants (Pontier et al., 2012), or in mammals, protein GW182 can, besides silencing, also modify the processing of mRNA (Elkayam et al., 2017).

Recently, another group and we showed that in *A. thaliana,* SPT6L colocalizes not only with Pol II but also with Pol V on chromatin, functioning in the RdDM pathway (Čermák et al., 2024; Liu et al., 2024). However, we also presented that the SPT6L protein is the most ancient AGO-hook domain-containing protein in any plant RNA polymerase complex, preceding even the emergence of Pol V (NRPE1). This suggests that the AGO-hook of SPT6L may have a specific function within the Pol II complex, affecting transcript processing or chromatin remodeling (Čermák et al., 2024).

Pol II-mediated mRNA transcription is a key process in gene expression with three highly regulated stages. It begins with initiation, during which the Pol II preinitiation complex assembles to recognize the proper transcription start site (TSS), open dsDNA, and initiate transcription. It is followed by productive elongation, during which the nascent transcript is synthesized. Finally, it concludes with transcription termination, leading to the stopping of the Pol II and the release of the final transcript (Schier and Taatjes, 2020). Notably, SPT6(L) has been implicated in all three of these transcriptional phases (Chen et al., 2019; Narain et al., 2021).

During transcription, multiple collateral processes occur, e.g., splicing, modulation of transcription dynamics, and epigenetic modifications of chromatin; the pre-mRNA can also be targeted by specific enzymes that deposit epitranscriptomic marks. One of the best researched epitranscriptomic marks so far is m^6^A methylation, which is deposited mainly in close relation to the poly-(A) site (PAS), leading to correct termination and influencing the stability of the mRNA product (Pontier et al., 2019; Zhang et al., 2024; Shao et al., 2025).

Recently, it has been shown that the RNA m^5^C mark is also an important mRNA posttranscriptional modification. The m^5^C mark is deposited on thousands of mRNAs, especially in their coding regions (David et al., 2017; Cui et al., 2017), with some research claiming that almost every transcript can be methylated (Song et al., 2018). Deposition of this mark to mRNA was shown to increase its stability (Warren et al., 2010), modulate translation activity (Cui et al., 2017; Tang et al., 2020), and influence mRNA nuclear retention (Yang et al., 2017; Pfaff et al., 2018). In plants, m^5^C exhibits tissue-specific enrichment during development and regulates key developmental transitions, including the maintenance of stem-cell proliferative capacity and root elongation (David et al., 2017). In *A. thaliana,* the most well-described RNA methyltransferase responsible for deposition of m^5^C marks is TRNA-SPECIFIC METHYLTRANSFERASE 4B (TRM4B), responsible for methylation of thousands of methylation sites with different distribution in plant tissues. While not highly abundant, the m^5^C mark in mRNA exhibits significant variability, indicative of its highly dynamic nature and rapid turnover (David et al., 2017; Cui et al., 2017).

Here, we show that the SPT6L protein, specifically through its AGO-hook domain, recruits AGO proteins to the Pol II complex and directs cytosine methylation of mRNAs. Furthermore, this methylation process is linked to transcription regulation, specifically by preventing premature termination.

## Results

### SPT6L interacts with AGO4 and AGO9 via the AGO-hook domain

To investigate the function of SPT6L’s AGO-hook, we generated gene constructs for expression of epitope-tagged full-length SPT6L (SPT6L-FLAG) and its version missing the AGO-hook (SPT6LΔAh-FLAG) under the control of the native promoter. These constructs were introduced into the *spt6l+/-* background. Both the full-length and truncated constructs fully complemented the lethal phenotype of the homozygous *spt6l-/-* null mutations, and plants exhibited normal vegetative growth with no obvious phenotypic alteration under standard cultivation conditions, consistent with a previous report (Figure 1A; Figure S1A; Chen et al., 2019). However, we observed a distinct flowering phenotype; plants complemented with the truncated SPT6LΔAh-FLAG version flowered approximately 4 days earlier than those complemented with the full-length SPT6L-FLAG version (Figure 1B; Figure S1B, C).

**Figure 1:**
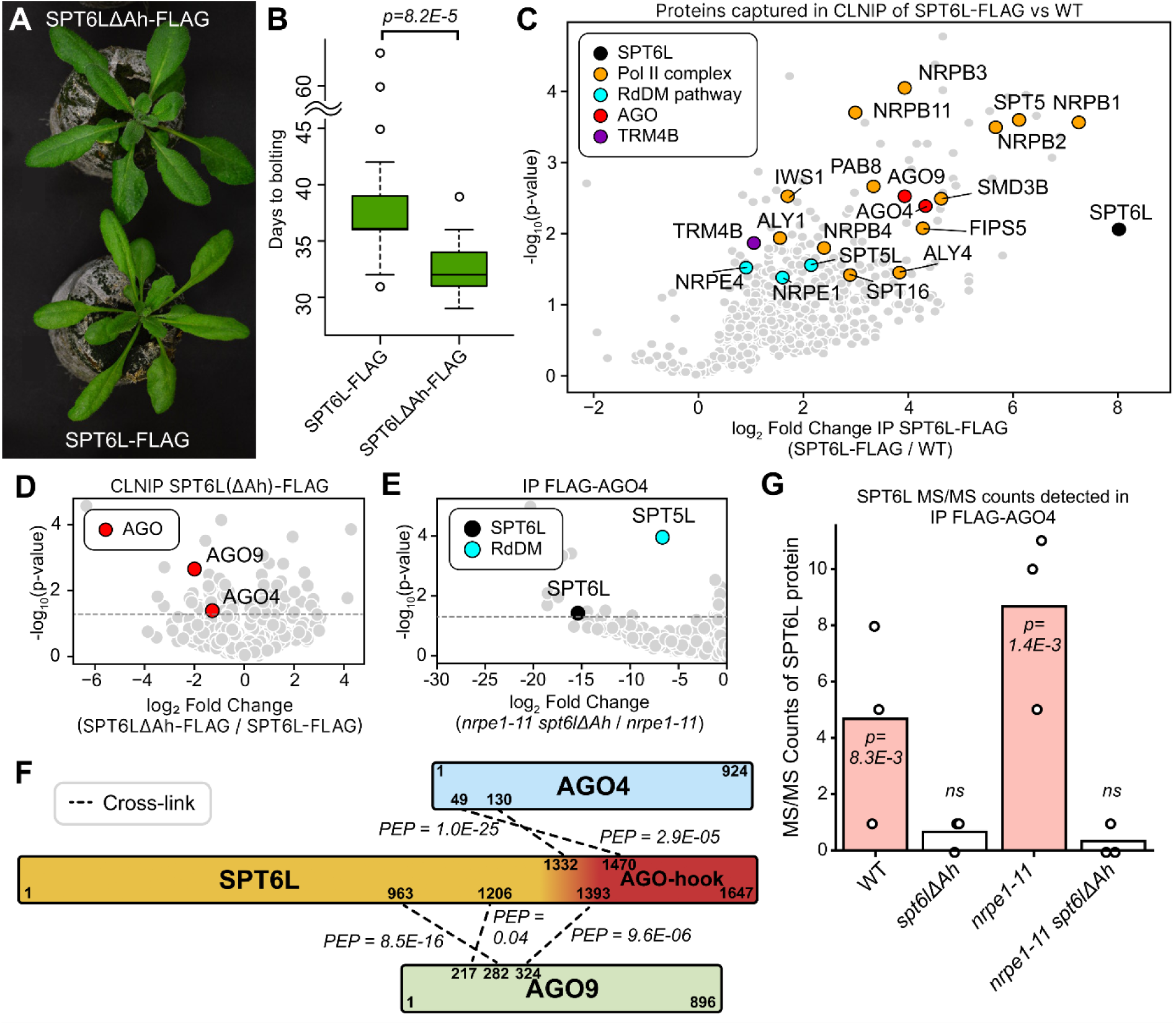
AGO-hook of SPT6L binds AGO proteins independently of Pol. **V** (A) Representative morphology of *spt6l-/-* null mutant plants complemented with either the full-length genomic SPT6L-FLAG construct (bottom) or the AGO-hook deletion variant SPT6LΔAh-FLAG (top). Both constructs rescued the *spt6l-/-* null mutation and supported vegetative growth. A comparison to WT Col-0 is available in Figure S1A. (B) Quantification of flowering time at long-day conditions measured as days to bolting in 30 plants per genotype. The SPT6LΔAh-FLAG plants flowered significantly earlier (approximately 4 days; two-tailed t-test p-value = 8.2E-5) compared to the full-length SPT6L-FLAG control. The representative morphology of flowering plants is shown in Figure S1B, and a comparison with WT is shown in Figure S1C. (C-E) Volcano plots showing differential enrichment of proteins captured in IP experiments. The x-axis indicates the difference between genotypes, and the y-axis represents the statistical significance. (C) Shows the difference between genotypes: SPT6L-FLAG / WT for CLNIP experiment. Key proteins enriched for SPT6L are color-coded: SPT6L (black), components of the Pol II complex (orange), AGO proteins (red), TRM4B (violet), and RdDM pathway components (cyan). We cropped the plot to show proteins enriched in SPT6L-FLAG (full data available in Supplementary Data S2). (D) Volcano plot showing differential enrichment of AGO4 and AGO9 proteins in CLNIP experiments for SPT6LΔAh-FLAG versus SPT6L-FLAG CLNIP samples comparison. (E) Differential enrichment of SPT6L protein in Co-IP experiments for comparison of FLAG-AGO4 *ago4-4 nrpe1-11 spt6lΔAh* versus FLAG-AGO4 *ago4-4 nrpe1-11* samples. The plot was cropped to show proteins enriched in FLAG-AGO4 *nrpe1-11* (full data available in Supplementary Data S2). The interaction between SPT6L and AGO4 is observed even in the *nrpe1-11* background and depends on the AGO-hook. (F) Graphical depiction of detected formaldehyde crosslinks in the CLNIP experiment between protein SPT6L (orange with AGO-hook denoted as red part) and either AGO4 or AGO9. For each crosslink, the PEP probability score and the sequence number of the paired amino acids are shown. Complete information about crosslinks is available in Supplementary Data S1. (G) Barplot of MS/MS counts of SPT6L protein captured in affinity assay for FLAG-AGO4 in various backgrounds, denoted on the x-axis - the “FLAG-AGO4 *ago4-4*” was omitted from the genotype name. The statistical comparison was done relative to the negative control (WT; Supplementary Data S2). The dots show MS/MS counts values for each biological replicate for the genotype.

Although the AGO-hook domain of SPT6L has been computationally predicted to serve for binding AGO proteins (Zielezinski and Karlowski, 2015), it has never been functionally verified. Therefore, we performed cross-linked nuclear immunoprecipitation (CLNIP) of both tagged SPT6L-FLAG and SPT6LΔAh-FLAG versions using nuclei isolated from young siliques treated with formaldehyde.

In pulldowns of full-length SPT6L-FLAG, we observed a significant enrichment of AGO4 and AGO9 compared to the control wild-type material (WT; Figure 1C, Supplementary Data S1). Importantly, this enrichment was significantly reduced in proteins co-immunoprecipitated with the truncated SPT6LΔAh-FLAG (Figure 1D, Supplementary Data S1). Furthermore, we detected cross-linked peptides between SPT6L and AGO4/9 in our data (Figure 1F, Supplementary Data S1). Consistent with previous studies, we detected numerous Pol II complex subunits and many other Pol II-associated proteins. Additionally, we detected protein SPT5L and two additional core subunits of the Pol V complex (Figure 1C, Supplementary Data S1), consistent with SPT6L’s dual role in both RNA polymerase complexes (Čermák et al., 2024; Liu et al., 2024).

To validate these interactions, we performed a reciprocal co-immunoprecipitation (Co-IP) assay with epitope-tagged AGO4. We utilized *ago4-4* mutants complemented with *pAGO4::FLAG-AGO4* construct (*FLAG-AGO4*; Wang and Axtell, 2017). We further generated a CRISPR/Cas9 mutant lacking the SPT6L AGO-hook (*spt6lΔAh*) and crossed it with *nrpe1-11* to generate a double mutant (*spt6lΔAh nrpe1-11*). We included *nrpe1-11* to deplete AGO4 interactome of proteins associated with its dominant role in the Pol V complex and RdDM, as this mutation renders the complex non-functional. This double mutant was subsequently crossed with *FLAG-AGO4* plants to introduce the epitope-tagged AGO4 and *ago4-4* mutation into all relevant mutant backgrounds. Subsequently, we performed native Co-IP experiments in total with five genotypes: WT Col-0 and four others on the *ago4-4 FLAG-AGO4* background: WT, *spt6lΔAh*, *nrpe1-11*, and *nrpe1-11 spt6lΔAh*. The Co-IP analysis confirmed efficient enrichment of AGO4 across all *FLAG-AGO4* genotypes compared with the WT control (Supplementary Data S2). As expected, known RdDM components, including the Pol V subunits NRPE1, NRPE2, and SPT5L, copurified with AGO4 in all genotypes except for the *nrpe1-11* - Pol V subunits mutant (Figure S1D, Supplementary Data S2). Importantly, we detected SPT6L in FLAG-AGO4 pulldown in WT but also in *nrpe1-11* mutant background, showing that the interaction between SPT6L and AGO4 is not exclusive to the Pol V complex (Figure 1G). Furthermore, the SPT6L signal was significantly depleted in the *spt6lΔAh nrpe1-11* background (Figure 1E, G). It is also worth noting that neither the interaction with SPT5L is lost in the *nrpe1-11* mutant.

In conclusion, our data show that SPT6L interacts with AGO4 and AGO9, and this interaction is dependent on the presence of the AGO-hook of SPT6L, but the AGO4-SPT6L interaction is independent of the presence of Pol V.

### SPT6L’s AGO-hook interaction with AGO4 guides RNA cytosine methylation

SPT6L serves as an interaction hub and multipurpose modulator of Pol II transcription, but the specific function of its AGO-hook domain, which anchors AGO proteins, remains unclear. Gene Ontology (GO) enrichment analysis of proteins interacting with SPT6L-FLAG revealed a significant overrepresentation of pathways involved in mRNA metabolism and processing (Figure S1E). We therefore leveraged Oxford Nanopore Technologies Direct RNA Sequencing (ONT DRS), which enabled us not only to evaluate gene expression changes but also to analyze the mRNAs from a wider perspective, including epitranscriptomic alterations in our AGO-hook CRISPR/Cas9 mutant *spt6lΔAh* and selected mutants with impaired sRNA production or function: *ago4-5, dcl234, nrpe1-11, nrpd1*. We chose *ago4-5* and not *ago9* mutant lines because *AGO9* is expressed only in generative organs, whereas *AGO4* is expressed almost ubiquitously (Havecker et al., 2010), allowing us to conduct the following experiments on seedlings.

First, we analyzed differential expression analysis for all mutants (Supplementary Data S3; Figure S2A). For *spt6lΔAh,* we identified 555 differentially expressed genes (DEGs) compared to WT, where most of them (431) were upregulated in the mutant (Figure S2B). These genes highly overlap with DEGs in *ago4-5* (n = 1 643; Figure S2A) and other mutants. Among these genes, we found significantly enriched genes in the GO categories “response to diverse stimuli” and “response to stress” (Figure S2C), for example, the *TAS3* gene, which controls development via posttranscriptional sRNA pathways. We also identified a few genes controlling the onset of flowering: AT5G47640 (NF-YB2), AT1G09570 (PHYA).

Next, we analysed changes in epitranscriptomic marks, and m^5^C stood out with interesting differences. Analysis of epitranscriptomic changes can be prone to error, leading to false-positive signals (Wiener and Schwartz, 2021). To overcome this, we applied stringent filters, retaining only highly and reproducibly methylated cytosines (see Methods). Altogether, we identified 13 430 highly and reproducibly methylated cytosines in the transcriptome of the analyzed *A. thaliana* genotypes (Supplementary Data S4). For these methylatable cytosines, we ran a differential methylation analysis, which revealed that differentially methylated cytosines (Dm^5^C) highly overlap across the analyzed genotypes (Figure 2A; Figure S3A, B). These changes do not seem to be due to differentially expressed RNA methyltransferases in the analysed mutants (Figure S2D).

**Figure 2:**
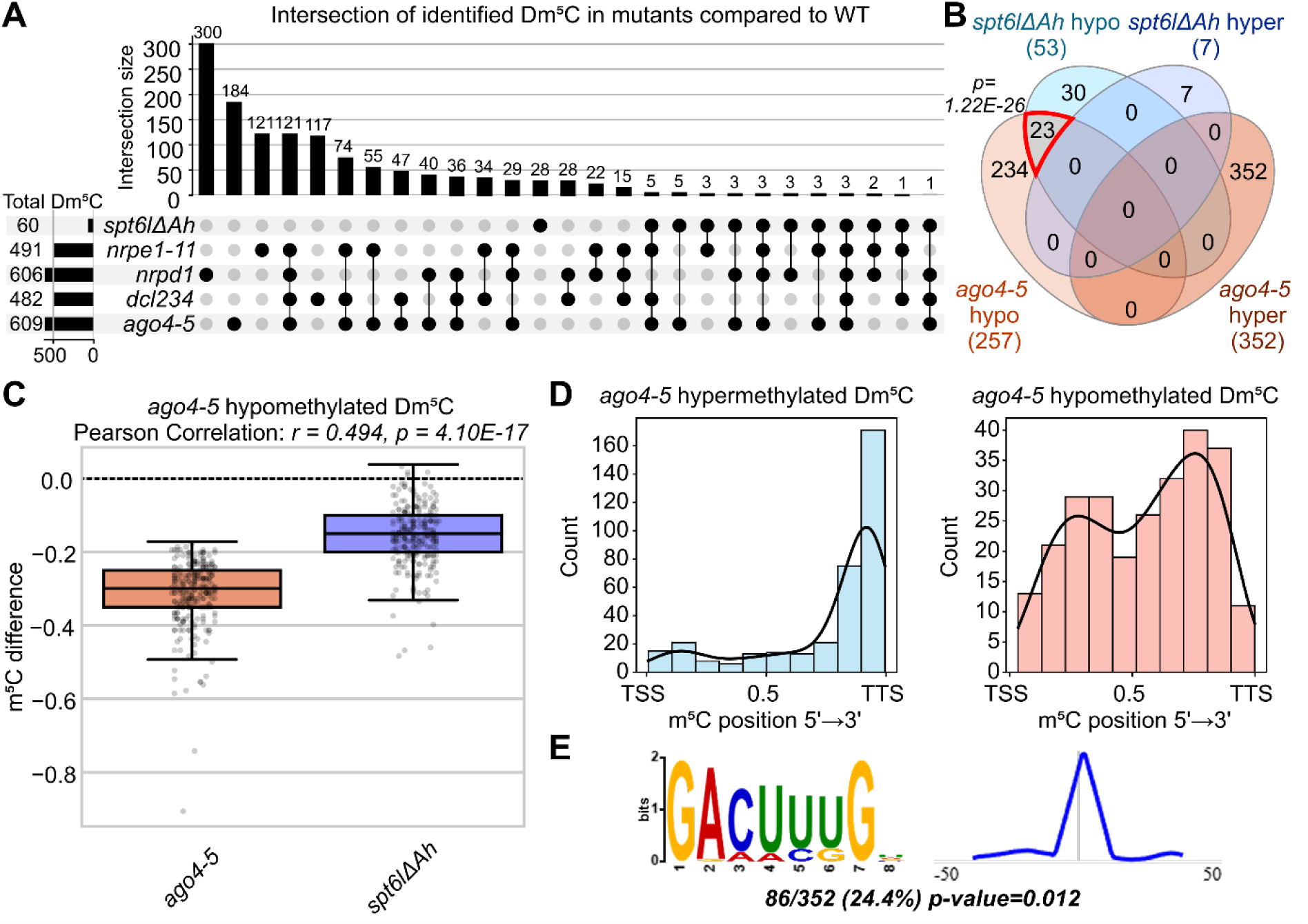
Characterization of m^5^C RNA methylation changes. (A) UpSet plot showing intersections of Dm^5^C identified in all studied mutants in comparison to WT. The UpSet plot, separated into hypo- and hypermethylated sites, is shown in Supplementary Figure S3A and S3B. Horizontal bars represent the total number of Dm^5^C for each genotype, while vertical bars show the number of sites shared between specific mutant intersections. (B) Venn diagram for Dm^5^C identified in *ago4-5* and *spt6lΔAh* in comparison to WT. Hypo- and hypermethylated sites were separated. The p-value for overlap of hypomethylated Dm^5^C was calculated using hypergeometric probability; p = 1.22E-26, and the representation factor is 23.9. (C) Boxplot of m^5^C difference showing m^5^C methylation change in either *spt6lΔAh* or *ago4-5* compared to WT on a set of hypomethylated Dm^5^C in *ago4-5* (n = 257). A Pearson correlation was used to test the association (r = 0.494, p = 4.10E-17). This positive correlation shows that the hypomethylation pattern is moving together in *ago4-5* and *spt6lΔAh*. (D) Histograms of Dm^5^C positions along their corresponding genes for hypermethylated Dm^5^C (n = 352; left) and hypomethylated (n = 257; right) in *ago4-5*. The hypomethylated Dm^5^C are mostly present in the 3’ end of the genes, whereas the hypomethylated Dm^5^C are distributed along the whole length of their genes. (E) The significant motif (p = 0.012) found using MEME-Chip for *ago4-5* hypermethylated Dm^5^C. The consensus sequence of this motif is GACUUUGW (left) with motif localization (right) centered on the hyper-Dm^5^C. The full results of the motif search are shown in Figures S3H and I.

When we analysed Dm^5^C in the *spt6lΔAh* mutant, we identified 60 sites, of which 53 were hypomethylated (Figure S3B; Supplementary Data S4). These hypomethylated cytosines significantly overlap with those significantly hypomethylated in *ago4-5* mutants (Figure 2B; Figure S3B), suggesting that the methylation of these cytosines might be part of a single pathway. That claim is also supported by the fact that 253 out of the 257 Dm^5^C hypomethylated in *ago4-5* showed lower methylation levels also in *spt6lΔAh* (Figure 2C). Also, other mutants show similar trends (Figure S3C, D). For those reasons, we continued with analysis focused on *ago4-5* Dm^5^C, which were more frequent but similarly affected in the *spt6lΔAh* mutant.

We stratified the identified sites based on methylation changes in the *ago4-5* mutant and observed distinct spatial distribution for hyper- and hypomethylated groups along the mRNA. While hypomethylated sites were almost evenly distributed, m^5^C hypermethylated in *ago4-5* were predominantly located near the 3’ end (Figure 2D). This distribution held strong also in the other mutants we analysed (Figure S3E). Next, we asked if mRNAs with Dm^5^C are differently expressed in our mutants. We did not find any significant correlation between the change in the level of mRNAs carrying Dm^5^C and the change in their methylation in any analyzed mutant (Figure S3F), which suggests that the RNA cytosine methylation is unlikely to be engaged in the regulation of mRNA levels or that the changes in methylation are a result of differential expression. Furthermore, we found that RNA methylation was not associated with DNA methylation in the CHH context, which is introduced via the RdDM pathway with the help of AGO4 (Figure S3H).

Since AGO4, together with Pol V, provides precise targeting of DNA methylation, we asked whether AGO4 may serve a similar function in the methylation of Pol II transcripts. Among the hits in the CLNIP interactors of SPT6L-FLAG, we captured the m^5^C RNA methyltransferase TRM4B and subunits of the THO/TREX complex (Figure 1C). The THO/TREX complex is an established component of the Pol II elongating complex (Grasser and Grasser, 2018) and is known to bind m^5^C on mRNAs, while TRM4B directly mediates methylation (David et al., 2017; Pfaff et al., 2018; Xu et al., 2024). We also captured some of these components in our FLAG-AGO4 Co-IP experiment (Supplementary Data S2). Together, this supports the idea that SPT6L bridges AGO proteins and the Pol II complex to recruit the TRM4B machinery and facilitate site-specific co-transcriptional methylation of those m^5^C sites, which were hypomethylated in the *ago4-5* or *spt6lΔAh* mutant backgrounds. In line with this supposed targeting of m^5^C by AGO4, which can be loaded with various sRNAs, we did not find any significant sequence motifs for cytosines hypomethylated in *ago4-5* using the MEME-ChIP tool, which would be indicative of the methylation targeting by RNA-binding proteins (Figure S3I). Conversely, within the *ago4-5* hypermethylated sites, we identified a significantly enriched consensus motif, GACUUUGW (p = 0.012; Figure 2E, Figure S3J), suggesting distinct mechanisms of targeting these hypermethylated Dm^5^C.

Among the genes with hypomethylated Dm^5^C in *ago4-5,* we found significantly enriched genes in the GO categories “gene expression” and “response to stress” (Figure S3G), and a few genes controlling the onset of flowering: AT3G04910 (WNK1), AT4G39400 (BRI1), and AT5G08330 (TCP21).

Furthermore, we confirmed that mRNA methylation levels were decreased in the *spt6lΔAh* and *ago4-5* mutants by sequencing of RNA immunoprecipitated with an antibody against m^5^C (Supplementary Data S4). We observed significant overlap of differentially methylated regions in *ago4-5* and *spt6lΔAh* when compared to WT, indicating that both mutated proteins participate in the same pathway (Figure S4A-C). The *ago4-5* mutation had a more pronounced effect on methylation levels, similar to that observed in the ONT experiment.

### AGO4:sRNA complex guides m^5^C methylation of specific mRNA in Pol II

The decrease in RNA methylation of specific cytosines was triggered not only by mutation of the AGO-hook domain of SPT6L and AGO4 but also by mutation in enzymes involved in sRNA production, such as DCLs, supporting the idea that sRNAs can guide targeting of mRNA methylation. Therefore, we first checked whether the AGO-hook of SPT6L is not responsible for generating sRNAs at these loci, rather than for guiding methylation. We performed sRNA-Seq in *spt6lΔAh* and WT and found no overlap between differential siRNA clusters and Dm^5^C loci (Supplementary Data S5). The 125 identified siRNA clusters differentially expressed in *spt6lΔAh* (Figure S5A) are probably connected to the function of SPT6L in RdDM, as DNA is methylated on these sRNA clusters and the methylation is lost in RdDM mutant *nrpe1-11* (Figure S5B). To determine whether AGO4 physically interacts with transcripts near the cytosines hypomethylated in *ago4-5*, we utilized publicly available data from AGO4 RIP-Seq (Au et al., 2017). We found specific enrichment of AGO4 right downstream of the hypomethylated Dm^5^C (Figure 3A). No such enrichment was found for control cytosines or hypermethylated Dm^5^C (Figure 3A, Figure S5C), and no AGO4 or some other RdDM components were detected in available ChIP-seq data on the template DNA around the hypo- and hypermethylated Dm^5^C (Figure S5D).

**Figure 3:**
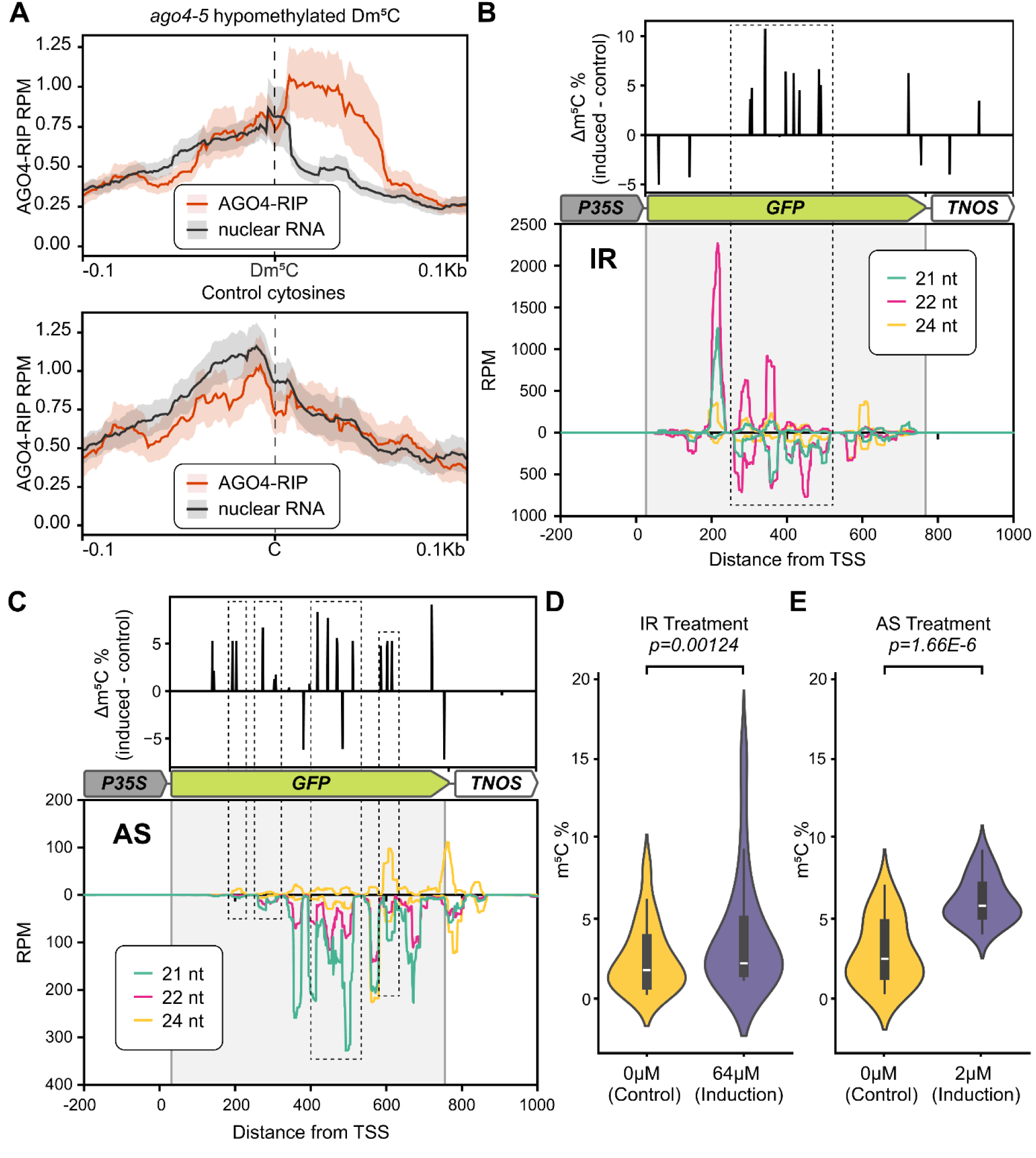
Role of AGO4:sRNA complex in guiding RNA m^5^C methylation. (A) Metaplot of AGO4 RIP experiment and the negative control (total nuclear RNA) from the GSE48617 dataset. The upper metaplot shows the ±100 bp region around *ago4-5* hypomethylated Dm^5^C (n = 257); the lower shows the same regions for control cytosines (random methylatable cytosines, n = 609). The lines show mean ± SEM as a shaded area. (B, C) Analysis of *GFP* mRNA methylation associated with the inverted repeat (*IR*; B) and antisense RNA (*AS*; C) silencing induction in tobacco BY-2 cells. The middle diagram shows the structure of the *P35S::GFP:TNOS* transgene. The lower panel displays sRNA coverage (coverage per million reads) mapped to the forward (top) and reverse (bottom) strands. Colors indicate different sRNA size classes: 21 nt (green), 22 nt (pink), and 24 nt (yellow). The upper panel shows the difference of m^5^C percentage between induced and control samples. The dashed box highlights the region showing high levels of specific reverse sRNAs correlating with increased RNA methylation in induced samples. (D, E) Violin plots comparing the distribution of the percentage of m^5^C difference between control and induced samples for the *IR* (D) and *AS* (E) lines. Statistical significance was tested using a two-tailed t-test.

To further test the hypothesis that RNA methylation, analogous to the RNA-directed DNA methylation pathway, can be guided by sRNA within the AGO4 complex, we utilized samples from our previous experiments with estradiol-inducible *GFP* silencing in *Nicotiana tabacum* BY-2 cell lines (Čermák et al., 2020). It allowed us to uncouple the developmental pleiotropy of *A. thaliana*, where m^5^C is known to differ across tissues (David et al., 2017), and to provide more direct proof of the concept of sRNA-guided methylation. In this system, cell lines carry a constitutively expressed GFP reporter, along with a second inducible cassette that triggers silencing by producing either antisense (*AS*) or inverted repeat (*IR*) *GFP* transcripts. Upon induction, these constructs give rise to double-stranded RNA, which is subsequently processed into sRNAs targeting the *GFP* transcript. To map the methylation landscape, we isolated total RNA before and after induction and performed ONT DRS. Although the level of *GFP* transcript was lower after induction due to RISC-mediated degradation, we could observe a distinct enrichment of m^5^C on the *GFP* transcripts in both induced lines compared to uninduced controls (Figure 3B, C). A quantitative analysis confirmed that the increase in methylation at methylatable cytosine sites was statistically significant for both systems: IR (p = 0.0012; Figure 3D) and AS (p = 1.66E-6; Figure 3E). Notably, the positional distribution of m^5^C closely mirrored the coverage profiles of the induced sRNAs for both *IR* and *AS* constructs (Figure 3B, C). This spatial correlation further supports that the specific sRNA distribution effectively predetermines the sites of RNA methylation.

### The m^5^C RNA methylation is connected with transcriptional pausing and release of Pol II

To assess the transcriptional consequences resulting from the absence of RNA m^5^C methylation, we crossed the *spt6lΔAh* mutant with *nrpb2-2* NRPB2-FLAG lines to perform plant Native Elongating Transcript sequencing (plaNET-Seq; Kindgren et al., 2020), providing a high-resolution map of transcribing Pol II positions via mapping of associated nascent transcripts. Our analysis revealed that the loss of the SPT6L AGO-hook led to global alterations in Pol II distribution. Specifically, we observed no difference in Pol II occupancy at the gene TSS (Figure 4A, B) and a decrease in the Pol II peak proximal to the poly-(A) site (PAS; Figure 4A, C), accompanied by increased Pol II occupancy in the gene body (Figure 4A). To investigate whether m^5^C presence or deposition influences these kinetic shifts and Pol II occurrence, we mapped plaNET-Seq signal relative to m^5^C. Since plaNET-Seq analysis indicated a global effect of the *spt6lΔAh* mutation on Pol II distribution, we filtered for all methylated cytosines that showed either a decrease or an increase in m^5^C methylation of at least 5% in the *spt6lΔAh* mutant compared to WT. The distribution of these sites along their respective genes mirrors the previously observed distribution of Dm^5^C (Figure 2D); cytosines hypermethylated in *spt6lΔAh* (m^5^C difference of > 5%; n = 1 316) occupy mostly 3’ end of the gene (Figure S6A), and hypomethylated cytosines (m^5^C difference of < −5%; n = 2 400) are more or less evenly distributed along the gene body (Figure S6B). Comparing these two cytosine sets, we observed a distinct accumulation of Pol II (plaNET signal), which peaked directly at cytosines with lower methylation and downstream of them. Importantly, the signal was significantly higher in the *spt6lΔAh* mutant (Figure 4D). This negative correlation strongly indicates that Pol II stalling is connected with these hypomethylated cytosines.

**Figure 4:**
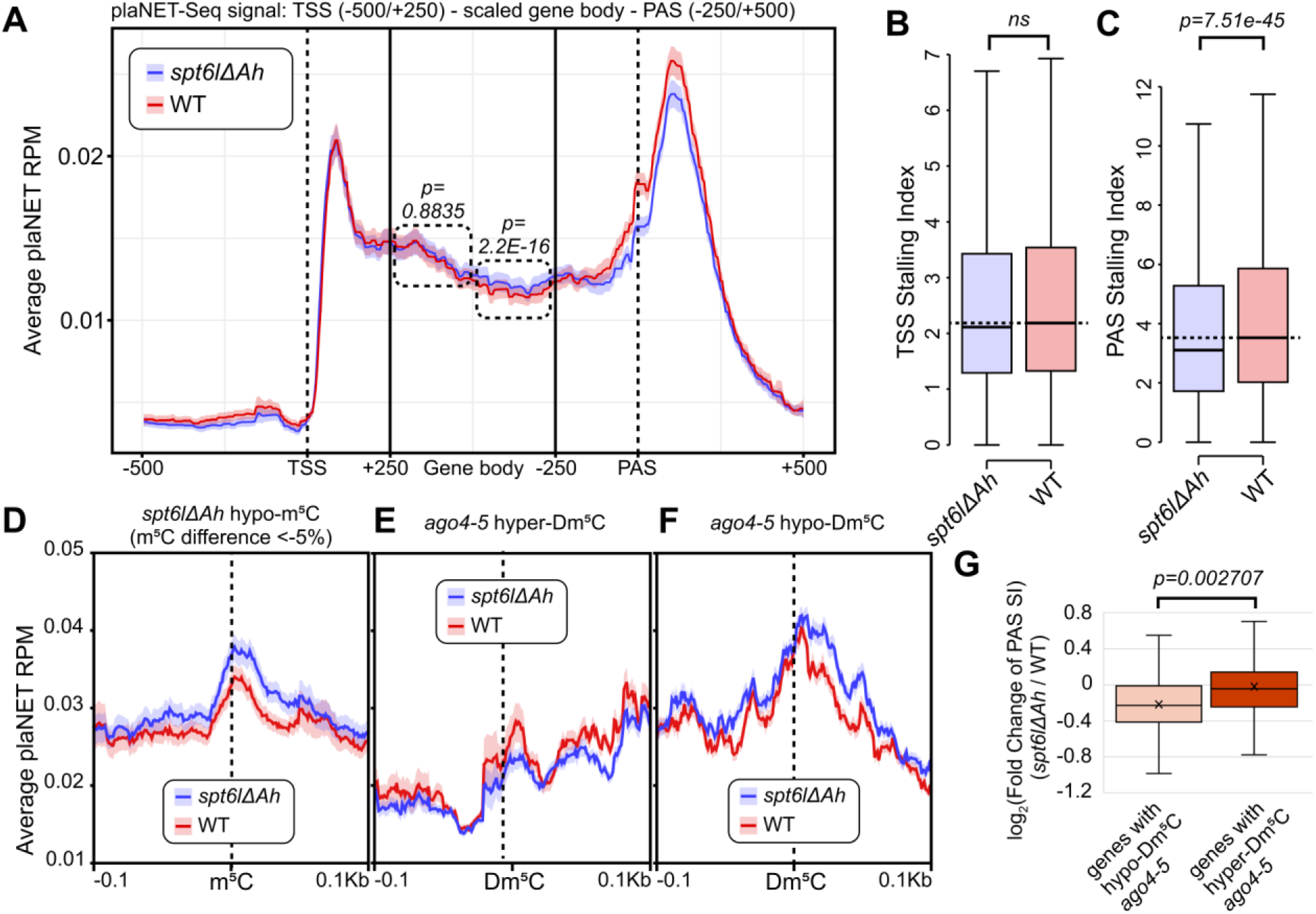
Loss of the SPT6L AGO-hook impairs Pol II processivity. (A) Metagene plot showing the average plaNET-seq signal ± 95% CI normalized to 1M tags in nuclear protein-coding genes plotted along all genes longer than 1kbp (n = 25 735) in WT (red) and *spt6lΔAh* (blue). The dashed rectangles indicate the half-regions within the gene body in which the difference in mean signal between the two conditions was quantified. The p-value was calculated using a Wilcoxon signed-rank test. The first half of the gene bodies shows a non-significant difference (p = 0.8835), whereas the second half shows a statistically significant difference (p = 2.2E-16) between WT and *spt6lΔAh* plaNET signals. (B, C) Boxplots comparing the TSS Stalling Index (TSS SI) between *spt6lΔAh* and WT (C; n = 27 353) or the PAS SI (D; n = 30 125). A two-sided Mann-Whitney U test was used to assess statistical significance. No statistically significant difference (ns) is observed around TSS, indicating transcription initiation is likely unaffected.  On the other hand, the *spt6lΔAh* mutant displays a significantly lower PAS SI (p = 7.51E-45), indicating a defect in elongation efficiency or termination. (D-F) Metaplots of the plaNET-seq signal were shown as mean ± SEM centered around specific cytosines. The three metaplots share the y-axis scale. (D) Metaplot of plaNET-seq signal flanking hypomethylated m^5^C, where the m^5^C difference of methylation change in *spt6lΔAh* in comparison to WT is < −5% (n = 2 400), Pol II density accumulates immediately downstream of *spt6lΔAh* hypomethylated m^5^C. (E, F) Metaplots of WT and *spt6lΔAh* plaNET-seq signal flanking Dm^5^C in *ago4-5* mutant; (E) hypermethylated sites and (F) hypomethylated sites. (G) Boxplots of log_2_ Fold Change of PAS SI for comparison of *spt6lΔAh* to WT for genes containing either hypo-(n = 257) or hypermethylated (n = 352) Dm^5^C in *ago4-5*. P-value was assessed by a two-sided t-test (p-value = 2.707E-3).

We did not observe similar Pol II stalling at hypermethylated cytosines or at control mRNA cytosines (Figure S6C, D).

Pol II stalling was not restricted to the cytosines that were hypomethylated in the *spt6lΔAh* mutant, we observed similar effect also at *ago4-5* hypomethylated Dm^5^C where Pol II became stalled downstream of *ago4-5* hypomethylated Dm^5^C sites. In contrast, at *ago4-5* hypermethylated Dm^5^C sites, Pol II showed reduced stalling (Figure 4E, F) as expected, because of their typical location closer to the gene terminus (Figure 2D) and mirroring the global trend in Pol II occupancy observed near the PAS site in the *spt6lΔAh* mutant (Figure 4B).

We further asked whether Pol II stalling at m^5^C sites with lower methylation within gene bodies can be linked to the global decrease in Pol II occupancy near the PAS site in the *spt6lΔAh* mutant (Figure 4C). Indeed, the decrease in Pol II occupancy at the PAS sites in *spt6lΔAh* was significantly lower in genes whose transcripts contained hypomethylated cytosines (m^5^C difference of < −5% in *spt6lΔAh*) compared to genes whose transcripts contained hypermethylated cytosines (Figure 4G).

To sum up, our data show that Pol II has a strong tendency to stall at the m^5^C sites. The stalling is further enhanced in the *spt6lΔAh* mutant, and is accompanied by decreased Pol II occupancy near the PAS site of the affected genes.

### m^5^C RNA methylation can drive transcription termination changes

Considering the lower Pol II occupancy near the PAS site in the *spt6lΔAh* mutant, we asked whether it may be caused by premature termination of Pol II transcription connected with stalling at hypomethylated m^5^C sites. To test this hypothesis, we used publicly available data for *trm4b* (Cui et al., 2017) and *ago4-2* mutants (Palomar et al., 2021), and compared them with Illumina steady-state RNA-Seq data from our *spt6lΔAh* mutants and respective WTs. The global coverage of the PAS region with Illumina reads for these mutants closely matches the signature of partial Pol II depletion at this region observed in the plaNET-Seq data (Figure S6F-H), supporting the idea of premature termination.

To analyze potential Pol II termination defects, we employed the APAtrap program (Ye et al., 2018). Comparing the *spt6lΔAh* mutant with the WT, we identified 710 differential termination events. Strikingly, 663 (93%) of these events corresponded to premature termination at proximal termination sites in the *spt6lΔAh* mutant, while only 47 (7%) preferentially terminated at distal sites (Figure 5A; Supplementary Data S6). Among genes with differential termination was AT2G21660 (GRP7), which is involved in the control of flowering time.

**Figure 5:**
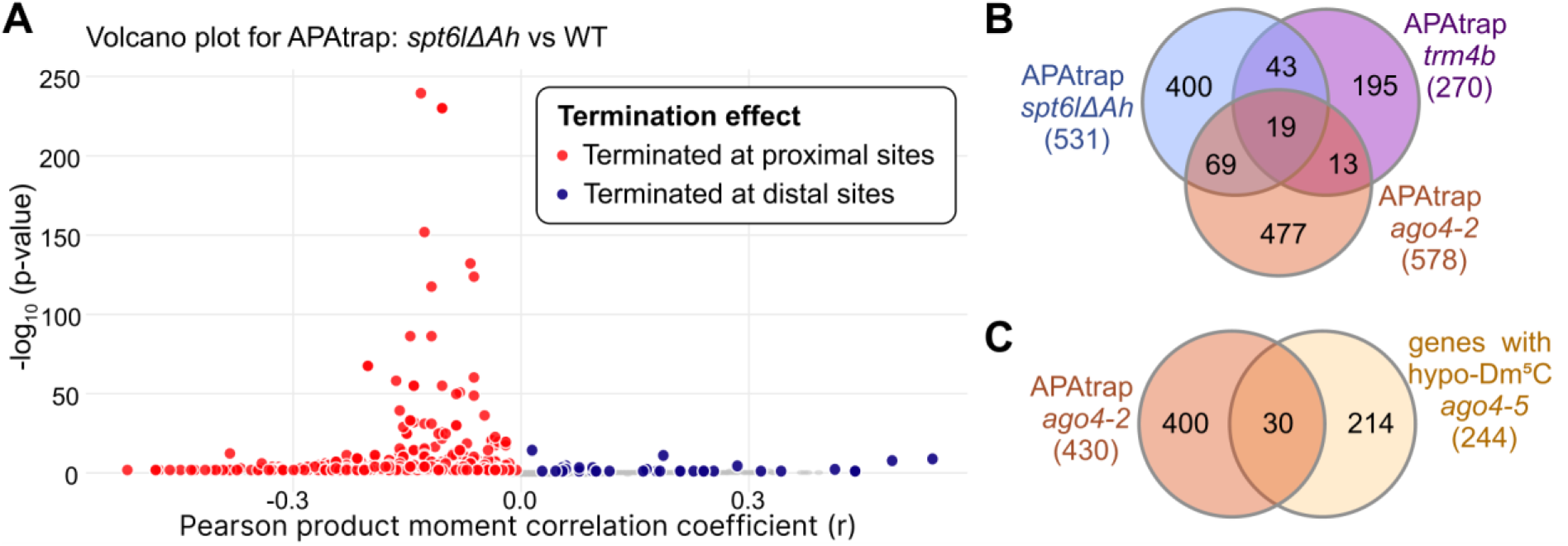
Pol II stalling in *spt6lΔAh* induces premature termination. (A) Volcano plot derived from APAtrap analysis comparing WT versus *spt6lΔAh* termination changes. The x-axis represents the Pearson product-moment correlation coefficient (r), and the y-axis represents the statistical significance. Red dots (r < 0) indicate genes showing premature termination, while blue dots (r > 0) indicate genes showing readthrough termination. (B, C) A Venn diagram illustrating the significant overlap of genes. Representation factors and hypergeometric probability p-values indicating highly significant overlaps between these datasets are available in Supplementary Data S6. (B) Overlaps for genes exhibiting differential termination (via APAtrap) in *spt6lΔAh*, *ago4-2*, and *trm4b* mutants. (C) Overlap of genes containing hypomethylated Dm^5^C sites in *ago4-5* genes with genes exhibiting differential termination (via APAtrap) in *ago4-2*.

Analysis of *the trm4b* mutant shows a similar but smaller effect: of 363 significant alternative termination events, 259 (71%) are prematurely terminated, and overlap significantly with genes prematurely terminated in *spt6lΔAh* (Figure 5B; Supplementary Data S6).

Furthermore, our analysis of the *ago4-2* mutant RNA-Seq data yielded comparable results, with 731 out of 773 (95%) events showing premature termination. When comparing these datasets of genes with diferrential termination events to one another, as well as to genes containing hypomethylated Dm^5^C cytosines in *ago4-5*, we observed significant overlaps (Figure 5B, C; Supplementary Data S6). Together, these results highlight the critical importance of AGO4, likely recruited to the Pol II complex via the AGO-hook of SPT6L, in guiding mRNA methylation to facilitate effective Pol II transcription elongation (Figure 6).

**Figure 6:**
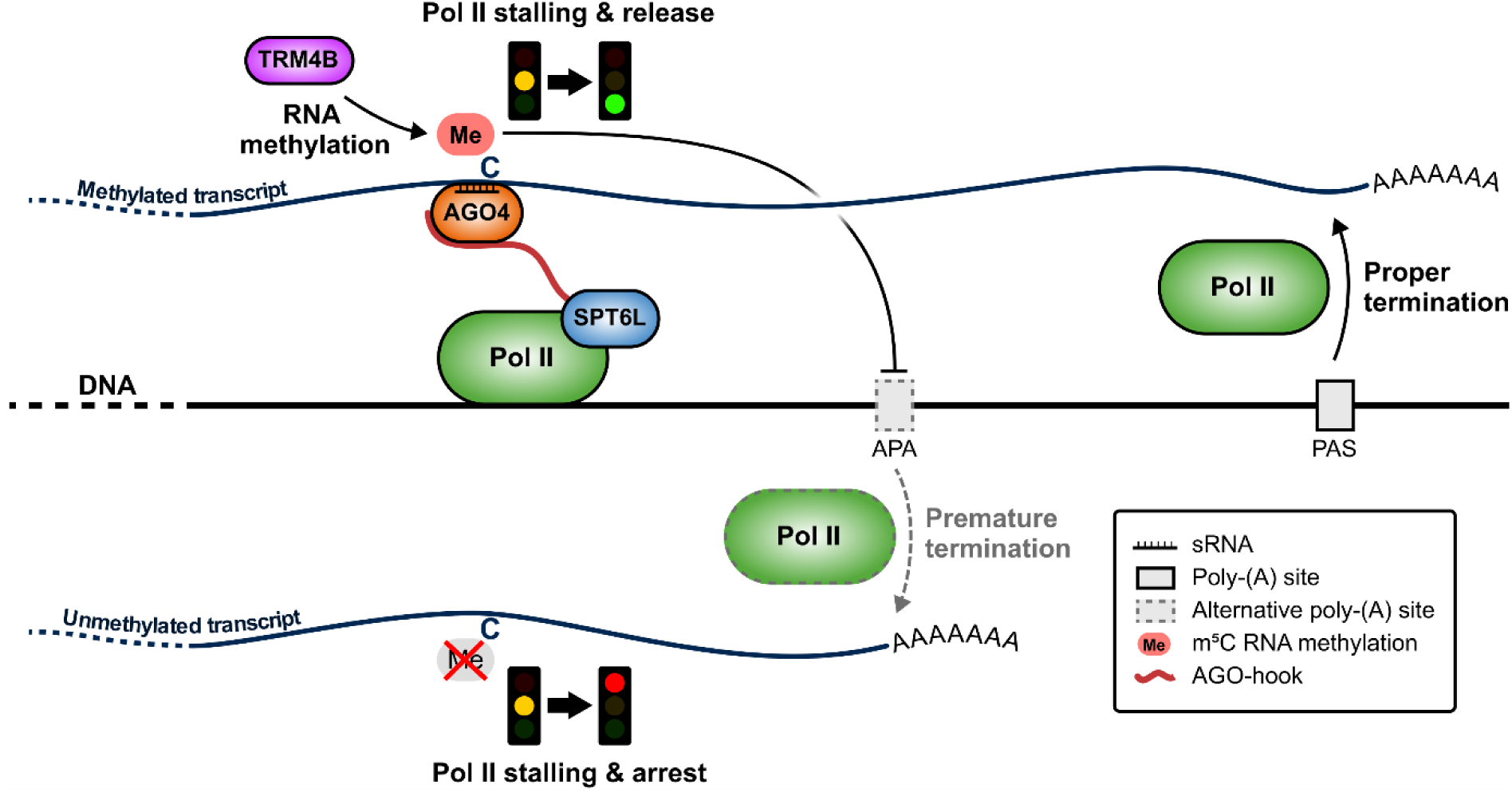
AGO4-directed mRNA m^5^C methylation prevents premature transcription termination in plants. Schematic illustration of the proposed mechanism linking sRNA-directed m^5^C mRNA methylation to the regulation of Pol II transcription elongation and termination in plants. The model contrasts two scenarios for nascent transcripts. During active transcription, SPT6L tethers sRNA-loaded AGO4 via its AGO-hook. This complex guides targeted m^5^C RNA methylation (probably via TRM4B RNA methyltransferase), which serves as a critical signal for Pol II to resume transcription after stalling and to continue to the proper polyadenylation site (PAS). On the other hand, if the mRNA is not methylated, Pol II arrests at the stall site, and transcription is terminated prematurely.

## Discussion

In this study, we identified SPT6L, an essential component of the Pol II complex, a histone chaperone, and a transcription elongation factor, as a new AGO-binding protein. The interaction with AGO4, and likely also with AGO9, is mediated by the C-terminal domain of SPT6L, which has previously been computationally predicted to be an AGO-hook domain (Karlowski et al., 2010; Zielezinski and Karlowski, 2015), but the interaction has not been experimentally verified until now. Previously, we showed that the AGO-hook of SPT6L has been the first one associated with RNA polymerase complexes to emerge in the plant lineage, preceding even the most ancient AGO-hook protein known at that time from the RdDM pathway - the Pol V subunit NRPE1 (Čermák et al., 2024). It suggests that the AGO-hook of SPT6L serves another function in the Pol II complex besides RdDM. Here, we show the interaction of SPT6L with RdDM pathway components, confirming that SPT6L is a physical component of the Pol V complex, as recent studies have shown (Čermák et al., 2024; Liu et al., 2024). Here, we also found that the absence of the AGO-hook affected the generation of sRNAs from 125 clusters linked to the RdDM pathway; despite this, Liu and colleagues did not observe significant changes in DNA methylation in a similar mutant (Liu et al., 2024). The missing effect on DNA methylation reported by Liu et al. (2024) also indicates that SPT6L’s AGO-hook might be functionally redundant with the other Pol V components with long AGO-hook domains, namely NRPE1 and SPT5L, whose partial redundancy was already demonstrated (Lahmy et al., 2016). However, the situation differs for the Pol II complex, where the AGO-hook of SPT6L should play a specific role. Here, we show that the interaction between AGO4 and SPT6L was preserved in the *nrpe1-11* background, which lacks the Pol V complex, suggesting that SPT6L tethers AGO4 (likely loaded with sRNA) to the Pol II complex. This is in line with previous studies demonstrating the presence of AGO4 in the Pol II complex (Zheng et al., 2009; Gao et al., 2010; Duan et al., 2015).

When focusing on the role of AGO4 in the Pol II complex, we found significant differences in m5C deposition in *spt6lΔAh, ago4-5,* and other sRNA pathway mutants. This suggests a new role for the sRNA machinery in regulating or navigating epitranscriptomic modifications, one that has not been described before.

Generally, epitranscriptomic modifications can be targeted by various mechanisms. In the case of m^6^A, targeting typically requires multiple layers of specificity, including sequence motifs recognized by the methyltransferase itself (e.g., the DRACH motif; Meyer et al., 2012) or by an accessory RNA-binding protein (e.g., RBM15; Patil et al., 2016). m^6^A deposition was also shown to be tightly linked to specific RNA structures (Spitale et al., 2015) or guided by the epigenetic histone modification H3K36me3 (Huang et al., 2019a). Targeting of m^5^C methylation is less well understood, but, in general, the known mechanisms are similar to those of m^6^A. For example, in mammals, some m^5^C was detected in the loop region of a stem-loop structure that harbored a 3′ TCCA motif (Huang et al., 2019b). One study of human cells also reported a positive correlation between m^5^C and m^6^A deposition (Li et al., 2017). Importantly, in humans, AGO proteins (hsAgo1-4), similar to our results, were shown to bind mRNA in the immediate vicinity of m^5^C sites, but without demonstrating causal relationship (Squires et al., 2012). Although hsSpt6 lacks an AGO-hook, other regulators such as GW182 utilize this domain to mediate repression and co-occupy transcripts with hsAgo1-4 (Hafner et al., 2010). Furthermore, Kim et al. (2024) recently demonstrated that RNA methylation can be deposited at sites where RNA forms double-stranded duplexes in mitochondria. Similarly, the RNA duplex formed between AGO-bound sRNA and its complementary transcripts could be targeted for methylation in the nucleus, as suggested by our results.

The hypothesis that sRNAs provide the sequence specificity for m^5^C RNA modification was strongly supported by our silencing inducible system based on *N. tabacum* BY-2 cells (Čermák et al., 2020), where the positional distribution of m^5^C well mirrored the coverage profile of the induced sRNAs (Figure 3B, C). This spatial correlation is a powerful argument for a “guide-and-modify” mechanism, in which the base-pairing of sRNA with the target mRNA could determine precisely where the methylation complex is deposited, and thus opens discussion for a possibly new pathway - RNA-directed RNA methylation. An analogous mechanism of snoRNA-guided posttranscriptional RNA modification is well established for the pseudouridinylation of tRNA and rRNA and may also target this modification within mRNAs, as recently documented (Nir et al., 2022). Our data also establishes a critical link between m^5^C RNA methylation and the kinetics of Pol II elongation. Using plaNET-Seq, we observed that loss of the SPT6L AGO-hook altered global Pol II distribution, characterized by increased signal within gene bodies on m^5^C sites and a reduced occupancy at PAS. Since no changes were detected at the TSS, this indicates a defect in processivity rather than initiation. Pol II specifically stalls downstream of cytosines that normally carry m^5^C marks, suggesting that methylation is mechanistically interconnected with the release of Pol II from internal pause sites. A similar function was described in yeast for the Spt6 protein. Here, the Spt6 protein recruits the m^6^A methyltransferase complex, which then triggers the release of Pol II following mRNA methylation at the promoter-proximal stalling site (Akhtar et al., 2021).

The stalling phenotype appears to drive premature transcription termination in all tested mutants affected in the presumed RNA-directed RNA methylation pathway, *spt6lΔAh*, *ago4,* and *trm4b*. However, the severity of the phenotype and the affected transcripts overlap only partially. These quantitative and qualitative differences suggest functional redundancy within the pathway; TRM4B may be redundant with any of the 8 other TRM4B paralogs found in *A. thaliana* (Wang et al., 2017). Similarly, AGO4 can be redundant with another AGO paralogue (Zhang et al., 2015), and the AGO-hook of SPT6L in the Pol II complex might be supplemented by the AGO-binding domain of the NERD protein (Pontier et al., 2019).

Taken together, we propose a novel “guide-and-modify” mechanism of RNA methylation, mediated by the tethering of sRNA-loaded AGO4 to the Pol II complex via the SPT6L AGO-hook. This targeted m^5^C deposition acts as a critical regulator of transcription, ensuring proper Pol II processivity and preventing premature transcript termination (Figure 6).

## Materials and Methods

### Plant Material

*pAGO4::FLAG-AGO4* expressed in the *ago4-4* (FLAG_216G02), *ago4-5* (WiscDsLox338A06), *dcl2/dcl3/dcl4* (SALK_064627, SALK_005512, GABI_160G05) hereafter referred to as *dcl234*, *nrpe1-11* (SALK_029919), *nrpd1* (SALK_143437), and *spt6l-2* (SAIL_569_E05) were obtained from the Eurasian Arabidopsis Stock Centre (uNASC). Seeds of *nrpb2-2* (SAIL_859B04) complemented by *pNRPB2::NRPB2-FLAG* were gifted to us by Dr. Sebastian Marquardt.

Fragments comprising the *pSPT6L::SPT6L(ΔAh)-StrepII-FLAG* construct were amplified using primers listed in Supplementary Data S7. Initially, the pGreen 0129 mCherry STOP plasmid (Hellens et al., 2000) was digested with enzymes EcoRI and SalI (excising the mCherry CDS), followed by the ligation of assembled oligonucleotides encoding the StrepII-FLAG tag. Then, the native promoter with the 5’ UTR part of SPT6L was amplified from *A. thaliana* Col-0 genomic DNA and inserted via SacI and EcoRI restriction sites. The SPT6L coding sequence was amplified from Col-0 cDNA in two parts (the second part was amplified using two different reverse primers, creating the full-length and AGO-hook-truncated versions of SPT6L). These two parts were assembled via the HindIII restriction site in the pDrive helper vector (QIAGEN). To create the final vector, the CDS was moved to the destination pGreen vector via BcuI and EcoRI restriction sites. This vector was transformed into *spt6l-2+/-*plants using a floral dip method. In the following generations, plants complementing this *spt6l-2-/-* knock-out mutation were selected.

For CRISPR/Cas9 deletion of the AGO-hook of SPT6L, the gRNAs were cloned into the pHSE401 vector following the protocol of Xing et al. (2014), using primers listed in Supplementary Data S7. The vector was transformed into Col-0 *Arabidopsis thaliana* plants, and in the next generation, a plant with the proper deletion between +6522-7428 bp of AT1G65440.1 was selected. The *pHSE401 T-DNA* was outcrossed, and the progeny plant without the T-DNA was once backcrossed to WT Col-0, and homozygous *spt6ΔAh* plants were selected. *A. thaliana* plants were grown ½ strength Murashige and Skoog media in a 16-hour light/8-hour dark photoperiod at a light intensity of 100 µmol · m^-2^ · s^-1^. Or in soil pellets with the same conditions.

The BY-2 cell line samples were from the study by Čermák et al. (2020), and all details on treatment and sample collection are described there.

### Crosslinked nuclear immunoprecipitation

CLNIP was done as previously described in Fang et al. (2020), with minor modifications. Briefly, 2 g of young siliques were harvested into 50 mL conical tubes and minced using scissors. Crosslinking was performed by vacuum infiltration for 12 min in 1% formaldehyde prepared in 1× PBS containing 0.01% Triton X-100. The reaction was quenched by adding glycine to a final concentration of 125 mM and incubating for 5 min under vacuum. Siliques were washed three times with 1× PBS, frozen in liquid nitrogen, and ground to a fine powder using a mortar and pestle.

Nuclei were isolated by incubating the ground tissue in 25 mL of Honda buffer (2.5% Ficoll 400, 5% Dextran T40, 0.4 M sucrose, 25 mM Tris-HCl, pH 7.4, 10 mM MgCl_2_, 10 µM β-mercaptoethanol, 0.5% Triton X-100, and 1× protease inhibitor cocktail Sigma cat. #P9599) for 10 min on ice with gentle shaking. The lysate was filtered through two layers of filter cloth, and the filtrate was centrifuged to pellet the nuclei.

The nuclear pellet was lysed in 200 μL of lysis buffer (50 mM Tris-HCl pH 8.0, 10 mM EDTA, 1% SDS, 1× protease inhibitor cocktail, Sigma cat. #P9599). This lysate was subsequently diluted to 2 mL with dilution buffer (1.1% Triton X-100, 1.2 mM EDTA, 16.7 mM Tris-HCl pH 8.0, 167 mM NaCl, 1× protease inhibitor cocktail Sigma cat. #P9599) and sonicated for 6 min at 50% power and a 50% duty cycle (Sonicator Bandelin UW2070). Protein complexes were immunoprecipitated using Anti-FLAG M2 Magnetic Beads (Millipore cat. #M8823) by incubating 20 μL of beads with 700 μL of the diluted nuclear extract for 1 h at 4°C. The beads were washed three times with 1× TBS. Proteomic analyses were performed at the OMICS Mass Spectrometry Core Facility at BIOCEV, Faculty of Science, Charles University.

### Native co-immunoprecipitation

Native co-immunoprecipitation (Co-IP) was performed as previously described by Liu et al. (2024). Briefly, 250 mg of young siliques were harvested, frozen in liquid nitrogen, and ground to a fine powder. The homogenized tissue was resuspended in 600 μL of lysis buffer (100 mM Tris-HCl pH 7.5, 300 mM NaCl, 2 mM EDTA, 1% Triton X-100, 10% glycerol, 1 mM PMSF, and 1× protease inhibitor cocktail, Sigma cat. #P9599) and incubated for 30 min on ice with gentle shaking. The lysate was then centrifuged at 18,000 × g for 10 min at 4°C, and the supernatant was collected. The cleared supernatant was incubated with 15 μL Anti-FLAG® M2 Magnetic Beads (Millipore cat. #M8823) for 2 h at 4°C on a rotary shaker. The beads were subsequently washed three times with a TBS buffer. Proteomic analyses were performed at the OMICS Mass Spectrometry Core Facility at BIOCEV, Faculty of Science, Charles University.

### Protein Digestion and nLC-MS/MS analysis

Protein digestion and proteomic analyses were performed at the OMICS Mass Spectrometry Core Facility at BIOCEV, Faculty of Science, Charles University. Briefly, proteins were digested overnight at 37 °C in 50 mM TEAB with 0.5 µg of trypsin. The supernatant was washed 3 times with ethyl acetate. Samples were acidified with TFA to a final concentration of 1%. Peptides were desalted using in-house-made stage tips packed with C18 disks (Affinisep, France) according to Rappsilber et al. (2007). nLC-MS/MS analysis was performed as follows: A Nano Reversed phase column (Ion Opticks, Aurora Ultimate TS 25×75 C18 UHPLC column) was used. Samples were loaded onto the trap column (C18 PepMap100, 5 μm particle size, 300 μm x 5 mm, Thermo Scientific) for 4 min at 18 μL/min. Eluting peptide cations were converted to gas-phase ions by electrospray ionization and analyzed on a Thermo Orbitrap Ascend (Thermo Scientific). Survey scans of peptide precursors from 350 to 1400 m/z were performed in Orbitrap at 120K resolution (at 200 m/z) with a 100 % ion count target. Tandem MS was performed by isolation at 1,6 Da with the quadrupole. Fragmentation spectra were acquired in ion traps with a Rapid scan rate. The MS2 ion count target was set to 150%, and the maximum injection time was 75 ms. Only those precursors with charge state 2-6 were sampled for MS2. The dynamic exclusion duration was set to 30 s, with a 10 ppm tolerance around the selected precursor and its isotopes. Monoisotopic precursor selection was turned on. Cycle time was set to 1.5 s.

### RNA and small RNA-seq

Total RNA was isolated from 12-day-old seedlings using TRIzol reagent (Invitrogen), following the manufacturer’s instructions. For steady-state RNA sequencing (RNA-seq) and small RNA sequencing (sRNA-seq), RNA samples were submitted to Novogene and BGI, respectively, in biological triplicates for library preparation and sequencing. Sequencing parameters for RNA-Seq were PE150 (∼40 million reads per sample), and for sRNA-Seq, SE50 (∼30 million reads per sample).

For ONT DRS, libraries for WT and *spt6lΔAh* were constructed in biological triplicate according to the manufacturer’s standard protocol for the Direct RNA Sequencing Kit (ONT cat. #SQK-RNA004) and sequenced for 48 h on a MinION RNA Flow Cell (ONT cat. #FLO-MIN004RA).

Libraries for *ago4-5, dcl234, nrpd1, nrpe1-11*, and BY-2 lines were constructed in biological triplicate using a custom multiplexing protocol for PromethION flow cells, derived from the WarpDemux protocol described by van der Toorn et al. (2025). Briefly, four custom-synthesized reverse transcription adapter (RTA) primers (Supplementary Data S7) were annealed to 250 ng of total RNA from each sample. Reverse transcription was performed according to the standard protocol, but scaled down to 1/4 volume. Barcoded samples were subsequently pooled and concentrated into a single sample during cleanup. The remainder of the library preparation followed the standard Nanopore DRS protocol. All libraries were sequenced using R10 flow cell chemistry (Supplementary Data S7).

### m^5^C-MeRIP

Total RNA was isolated from 100 mg of 12-day-old seedlings using TRIzol (Invitrogen) according to the manufacturer’s instructions. 40 μg of the total RNA was treated with TURBO DNAse (Invitrogen cat. #AM2238), and poly-(A) mRNA was isolated using NEBNext® High Input Poly(A) mRNA Isolation Module (NEB #E3370S). The poly-(A) enriched RNA was fragmented by adding a 2x Alkaline RNA fragmentation buffer (2 mM EDTA, 10 mM Na_2_CO_3_, 90 mM NaHCO_3_) and heating to 95°C for 11 minutes. Fragmented RNA was precipitated by ethanol extraction with the addition of Glycoblue (Invitrogen cat. #AM9515) and diluted to 430 μL. The RNA was mixed with 50 μL of 10x MeRIP buffer (100mM Na-Phosphate buffer pH 7.0, 1.4M NaCl, 0.5% Triton-X 100), 5 μg of α-m^5^C (Proteintech cat. #68301-1-Ig) or IgG (Proteintech cat. #65128-1-Ig) antibody, and 10 μL of RNAse inhibitors (NEB cat. #M0314S). The RNA was incubated with antibodies for 16 h. Then, 40 μL of pre-equilibrated protein-A magnetic beads (Invitrogen cat. #2791319) were added and incubated for 2 h at 4 °C. After that, the beads were washed three times using a 1× MeRIP buffer. 200 μL Proteinase K digestion buffer (50 mM Tris-HCl, pH 8.0, 10mM EDTA, 0.5% SDS) was added along with 3.5 μL RNase grade Proteinase K (NEB cat. #P8107S) and incubated at 50°C for 1 h on a shaker. The immunoprecipitated RNA was isolated using TRIzol (Invitrogen) with the addition of Glycoblue (Invitrogen cat. #AM9515), diluted to 12 μL or RNAse-free water, and sent for sequencing to Novogene (∼30 million PE150 reads on NovaSeq X).

### plaNET-Seq

plaNET-Seq was performed as previously described by Kindgren et al. (2020), with some modifications. Nascent RNA was isolated from 3g of 12-day-old seedlings. Following the initial isolation steps, the anti-FLAG antibody-conjugated beads (Millipore cat. #M8823) were washed six times. RNA was subsequently isolated directly from the beads using TRIzol reagent (Invitrogen) with the addition of Glycoblue (Invitrogen cat. #AM9515), following the manufacturer’s instructions. For library construction, 100 ng of nascent RNA was concentrated to a final volume of 5 µL using AMPure XP beads (Beckman Coulter cat. #A63987).

Due to the discontinuation of the kit used in previous protocols, the NEXTFLEX Small RNA Sequencing Kit V4 (Revvity cat. #NOVA-5132-32) was employed instead. Following the initial 3′ adapter ligation and heat inactivation, the RNA was fragmented by incubation in an equal volume of 2× alkaline fragmentation solution (2 mM EDTA, 10 mM Na_2_CO_3_, 90 mM NaHCO_3_) at 95°C for 1 min, a duration optimized to yield approximately 100 nt fragments. The fragmented RNA was purified using RNAClean XP beads (Beckman Coulter cat. #A63987), treated with T4 Polynucleotide Kinase (NEB cat. #M0201S) in T4 DNA ligase buffer (NEB cat. #B0202S), and purified again. The library preparation protocol was then reinitiated from the beginning according to the manufacturer’s instructions, omitting the 3′ RNA adapter addition. Final libraries were submitted to Novogene for deep sequencing (∼100 million paired-end 150 bp reads per sample) on the NovaSeq X Plus platform.

### Bioinformatic analysis

#### MS/MS data analysis

Raw MS/MS data were processed using MaxQuant (v2.4.2.0) for peptide identification. Native IP data were analyzed using default parameters. For CLNIP data, parameters were adjusted to allow for up to 3 missed cleavages and a maximum peptide mass of 6000 Da. All searches were performed against the *A. thaliana* reference proteome (UniProt UP000006548).

Statistical analyses were conducted using Perseus (v2.0.10.0), and data visualization was performed using custom scripts. To avoid bias when comparing full-length SPT6L with the deletion mutant (SPT6LΔAh), truncated SPT6L annotations were used for quantification in both samples. For crosslinking analysis, MaxQuant was run using non-cleavable formaldehyde crosslinking settings, specifically targeting only SPT6L (UniProt: A8MS85), AGO4 (Q9ZVD5), and AGO9 (Q84VQ0).

#### ONT DRS data analysis

Multiplexed libraries were demultiplexed using WarpDemux (v1.0.0) with the WDX4b_rna004_v1 model. Reads were assigned to specific barcodes and sorted into separate files using the custom script match_barcodes_to_reads.py, available in the associated Codeberg repository. Subsequent processing followed the standard pipeline used for non-multiplexed libraries.

Basecalling was performed using Dorado v1.3.0 with the super-accuracy (sup) model (rna004_130bps_sup@v5.1.0), configured to detect m^5^C RNA modification and estimate poly(A) tail lengths. Reads were then aligned to the *A. thaliana* TAIR10 reference genome using the built-in Dorado aligner.

For differential expression analysis, gene counts were generated with featureCounts v2.0.6 using the - L parameter for long reads, and statistical analysis was performed with DESeq2. RNA modification analysis was conducted using modkit v0.4.4 (pileup and dmr subcommands) with default parameters.

For each genotype, the high-confidence m^5^C site must fulfill these criteria: the overall count of reads with that specific m^5^C modified is larger than 2 reads + 2 RPM, the overall ratio of reads with that specific cytosine methylated is at least 20%, and the total number of reads on that specific cytosine is larger than 5 reads + 10 RPM. The cytosines identified this way in all genotypes were pooled into a methylatable cytosine database (Supplementary Data S4). For these cytosines, differential methylation analysis was performed, and the resulting MAP p-value was corrected for multiple testing using the Bonferroni-Hochberg method. We used the modkit effect size as the m^5^C difference, which is a change in methylation level ranging from −100% (hypomethylation) to 100% (hypermethylation). As a control cytosine set, we used 609 cytosines (the number of Dm^5^C in *ago4-5*), shuffled across methylatable cytosines with bedtools shuffle.

### m^5^C-MeRIP data analysis

The raw reads were aligned to the Araport11 transcriptome using bowtie2 (v2.4.2) in paired-end mode with the --sensitive-local option. Peaks were called using macs2 callpeak (v2.2.7.1) with parameters: --gsize ‘45000000’ --format BAMPE --keep-dup ‘auto’ --qvalue ‘0.05’. Reads were filtered using samtools (v1.14) to remove duplicates and keep only properly paired reads. Deferentially enriched peaks were then identified using DiffBind (v3.16.0) with method DBA_DESEQ2. Differentially enriched peaks with FDR < 0.05 were considered significant.

### plaNET-Seq data analysis

plaNET-Seq data were processed following the pipeline described by Kindgren et al. (2020), with minor modifications. Briefly, raw reads were adapter-trimmed using Cutadapt v5.1 with -m 15. Mapping to the *A. thaliana* TAIR10 genome was performed using STAR v2.7.11b with parameters optimized for short nascent reads: --outFilterScoreMinOverLread 0.1, --outFilterMatchNminOverLread 0.1, -- outFilterMatchNmin 0, --alignIntronMax 5000, --outFilterMultimapNmax 5, and -- outSAMmultNmax 1.

Following alignment, only R2 reads were used, and reads mapping to specific unwanted loci (e.g., rRNAs, tRNAs) were removed using bedtools intersect, as described in the Kindgren et al. (2020) protocol. Finally, PCR duplicates were removed using samtools markdup.

### Illumina RNA-Seq and sRNA-Seq data analysis

Steady-state RNA-Seq data were trimmed using Cutadapt v5.1 with parameters -m 15 -M 150 to remove adapters and short reads. The trimmed reads were aligned to the *A. thaliana* TAIR10 reference genome using STAR v2.7.11b, and reads with a mapping quality (MAPQ) score < 10 were discarded. Alternative polyadenylation (APA) analysis was performed using APAtrap with the following specific parameters: identifyDistal3UTR (-w 50 -e 2500 -p 0.9 -c 0.1), predictAPA (-a 50 -c 20 -d 0.1), and deAPA (least_qualified_num_in_group1 2, least_qualified_num_in_group2 2, coverage_cutoff 20). sRNA data were mapped and analyzed using ShortStack v4.0.4 (Axtell, 2013). The analysis was configured with the parameters --pad 100, --dicermin 20, --dicermax 25, and --nohp.

### Metagene plots

Metaplots were created using deeptools v3.5.6 using computeMatrix and plotProfile commands. We used the mean with the standard error of the mean for plotting with 1 bp bins.

The metaplot for Figure 4A was plotted using a custom script adapted from Kindgren et al. (2020) and is available in the Codeberg repository for this study.

### Gene Ontology

Gene Ontology (GO) analysis and plots were generated using the ShinyGO web application (v0.85.1; Ge et al. (2020).

## Data availability

The scripts required to reproduce our results are available at the Codeberg repository: https://codeberg.org/kaspartom/Kaspar_et_al_2026.git.

Complete sequencing data sets generated from this publication have been deposited in NCBI’s Gene Expression Omnibus and are accessible through accession number GSE324578.

Complete proteomic data were deposited in the PRIDE database under the accession number PXD075512.

## Author contributions

T.K., V.C, K.A., and L.F. conceived and designed the experiments; K.A. helped perform and managed T.K. with plaNET-Seq, and initial ONT DRS; V.C. generated all the constructs and initial plant genotypes, performed the phenotype analysis and m^5^C-RIP-Seq analysis; T.K. performed all proteomic and sequencing experiments and analysed all data for CLNIP, Co-IP, RNA-Seq, sRNA-Seq, ONT DRS, and plaNET-Seq experiments; T.K., V.C, K.A., and L.F. wrote the paper. All authors read and approved the final article.

## Acknowledgments

Computational resources were provided by the e-INFRA CZ project (no. 90254), supported by the Ministry of Education, Youth and Sports of the Czech Republic.

Proteomic analyses were performed at the OMICS Mass Spectrometry Core Facility at BIOCEV, Faculty of Science, Charles University.

We thank Dr. Sebastian Marquardt for providing the NRPB2-FLAG seeds used in this study. We thank Dr. Peter Kindgren and Dr. María Luz Annacondia López for their consultation and troubleshooting on plaNET-Seq.

