## Supplementary Figures and Legends for "Transcription elongation factor SPT6L recruits ARGONAUTE to guide mRNA cytosine methylation preventing premature termination in plants"

**Supplementary Data**

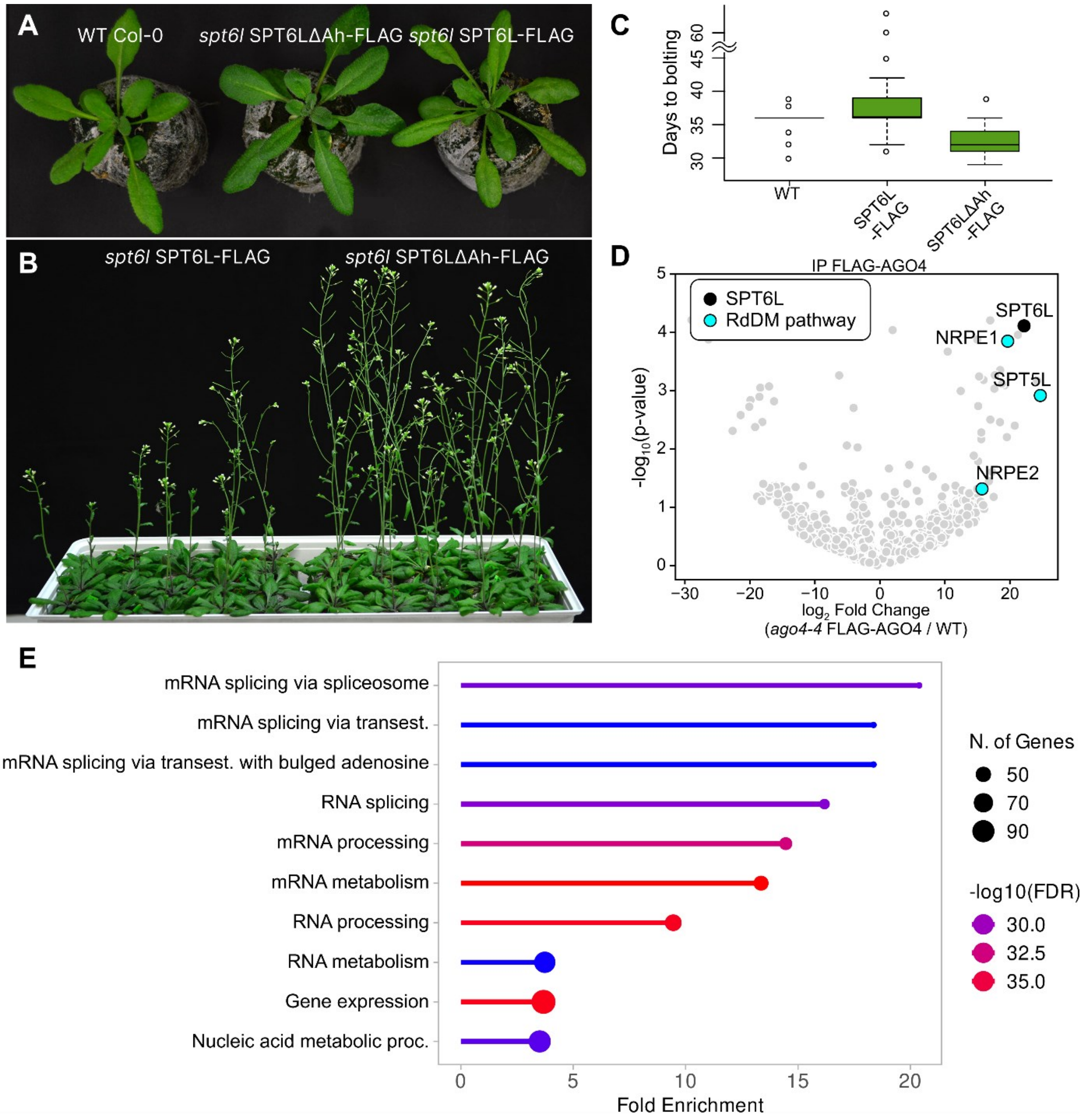


**Figure S1: AGO-hook of SPT6L interacts with AGO4, and mutant plants have a distinctive flowering phenotype**

(A) Representative morphology of WT Col-0 plants and *spt6l-/-* null mutant plants complemented with either the full-length genomic SPT6L-FLAG construct (pSPT6L::SPT6L-StrepII-FLAG) or the AGO-hook deletion mutant SPT6LΔAh-FLAG (pSPT6LΔAh::SPT6LΔAh-StrepII-FLAG).

(B) Representative flowering phenotype of *spt6l* null plants complemented with either full-length genomic SPT6L-FLAG (left) or the AGO-hook deletion mutant SPT6LΔAh-

FLAG (right). The *SPT6LΔAh-FLAG* plants exhibit an early-flowering phenotype, as evidenced by more advanced inflorescence development compared to the control. The plants were grown in randomized positions on the tray and reorganized solely for this picture.

(C) Quantification of flowering time measured as days to bolting in 30 plants per genotype. The graph is related to Figure 1B.

(D) Volcano plot showing enrichment of proteins co-immunoprecipitated with FLAG-AGO4 compared to a WT negative control. The x-axis represents the $log_2$ fold change enrichment of FLAG-AGO *ago4-4* vs WT Col-0 samples, and the y-axis represents statistical significance. Black dots highlight SPT6L, and cyan dots highlight known RdDM pathway components (NRPE1, NRPE2, SPT5L) that interact with AGO4.

(E) Lollipop plot of Gene Ontology (GO) enrichment of proteins enriched in the SPT6L-FLAG compared to WT in the CLNIP experiment.

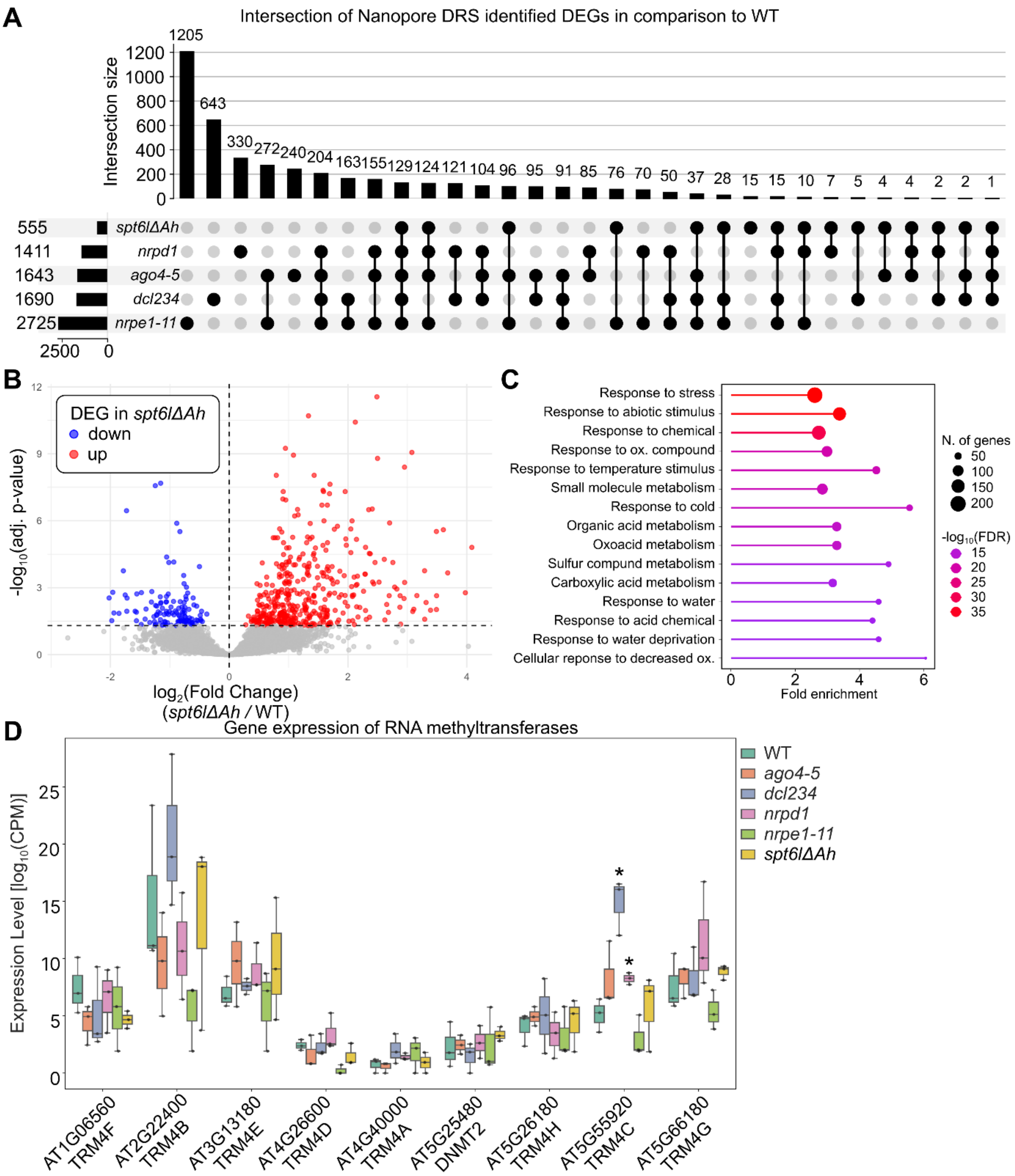


**Figure S2: Mutation of SPT6L AGO-hook influences stress gene expression**

(A) UpSet plot of differentially expressed genes detected by DESeq2 for mutants *ago4-5*, *nrpe1-11*, *spt6lΔAh*, *nrpd1*, *dcl234* in comparison to WT. Full information on the DEGs is available in Supplementary Data S3. Horizontal bars represent the total number of DEGs for each genotype, while vertical bars show the number of sites shared between specific mutant intersections.

(B) Volcano plot for DEGs in *spt6lΔAh* compared to WT. The upregulated genes are denoted in red, and the downregulated genes are denoted in blue. The x-axis shows $log_2$ Fold Change for the genes computed by DeSeq2, and the y-axis shows statistical significance.

(C) Lollipop plot of GO enrichment for *spt6lΔAh* DEGs.

(D) Boxplots of expression levels in normalized counts of various $m^5C$ probable methyltransferases: AT1G06560, AT2G22400 (TRM4B), AT4G26600, AT4G40000, AT3G13180, AT5G25480, AT5G26180, AT5G66180, AT5G55920, and AT5G66180. (*) denotes significant changes in expression identified by DESeq2 in relevant comparison to WT, the only significantly changed methyltransferase gene is AT5G55920 (TRM4C) in *dcl234* and *nrpd1*.

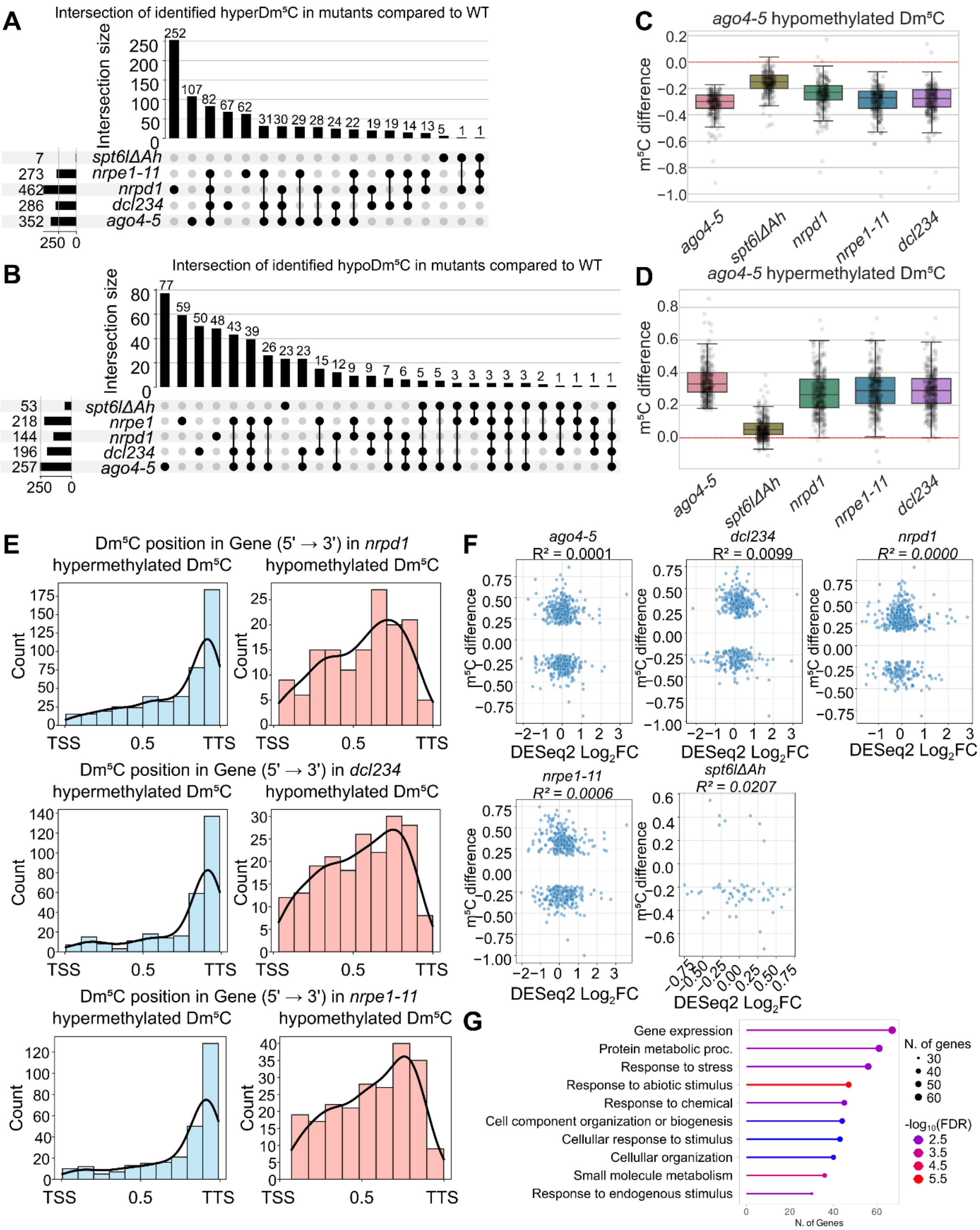
A
Intersection of identified hyperDm⁵C in mutants compared to WT
Intersection size
252
107
82
67
62
31
30
29
28
24
22
19
19
14
13
5
1
1
7 spt6lΔAh
273 nrpe1-11
462 nrpd1
286 dcl234
352 ago4-5
B
Intersection of identified hypoDm⁵C in mutants compared to WT
Intersection size
53 spt6lΔAh
218 nrpe1
144 nrpd1
196 dcl234
257 ago4-5
C
ago4-5 hypomethylated Dm⁵C
m⁵C difference
ago4-5
spt6lΔAh
nrpd1
nrpe1-11
dcl234
D
ago4-5 hypermethylated Dm⁵C
m⁵C difference
E
Dm⁵C position in Gene (5' → 3') in nrpd1
hypermethylated Dm⁵C
hypomethylated Dm⁵C
Dm⁵C position in Gene (5' → 3') in dcl234
Dm⁵C position in Gene (5' → 3') in nrpe1-11
Count
TSS
0.5
TTS
F
ago4-5
R² = 0.0001
dcl234
R² = 0.0099
nrpd1
R² = 0.0000
nrpe1-11
R² = 0.0006
spt6lΔAh
R² = 0.0207
m⁵C difference
DESeq2 Log₂FC
G
Gene expression
Protein metabolic proc.
Response to stress
Response to abiotic stimulus
Response to chemical
Cell component organization or biogenesis
Cellular response to stimulus
Cellular organization
Small molecule metabolism
Response to endogenous stimulus
N. of Genes
N. of genes
30
40
50
60
-log₁₀(FDR)
2.5
3.5
4.5
5.5

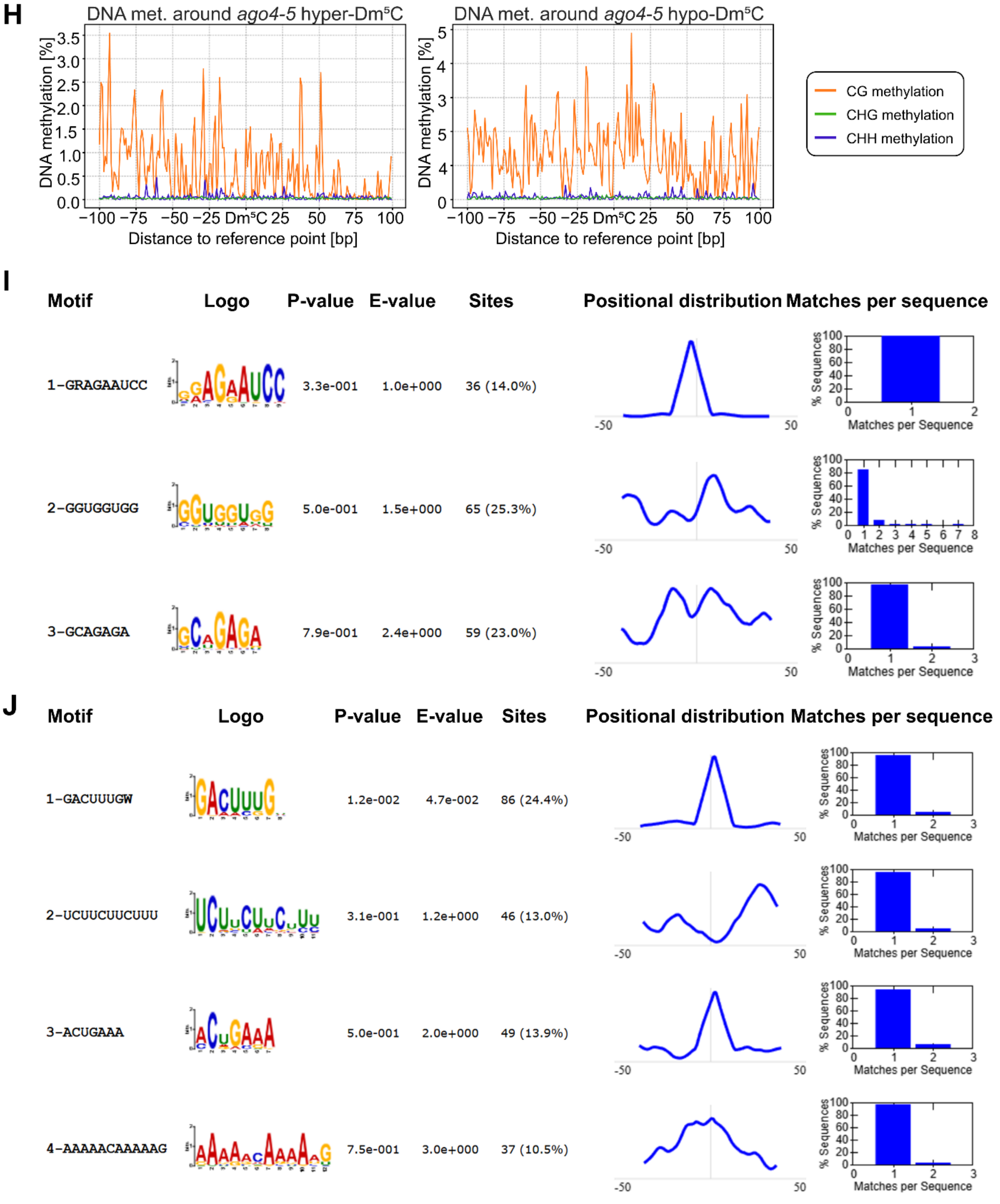


**Figure S3: Mutation of sRNA pathway components changes the m$^5$C deposition in mRNA**

(A, B) UpSet plots displaying the intersection of hypermethylated (A) and hypomethylated (B) Dm$^5$C identified in *spt6lΔAh*, *nrpe1-11*, *nrpd1*, *dcl234*, and *ago4-5* mutants in comparison to WT. Horizontal bars represent the total number of Dm$^5$C for each genotype, while vertical bars show the number of sites shared between specific intersections of mutants. Higher methylation, especially in *nrpd1* and *dcl234* mutants, could be connected to the higher expression of RNA methyltransferases (Figure S2D).

(C, D) Box plots showing the methylation $m^5C$ difference (change in methylation level from -1 hypomethylation to 1 hypermethylation) in the indicated genotypes at $Dm^5C$ sites specifically identified as hypomethylated (C) or hypermethylated (D) in the *ago4-5* mutant. The red dashed line represents zero change. Note that *spt6lΔAh* exhibits a decrease in methylation at *ago4-5* hypomethylated sites but lacks the large hypermethylation signature seen in other RdDM mutants.

(E) Histograms of distribution of hypermethylated (blue) and hypomethylated (red) $Dm^5C$ along their corresponding genes for *nrpd1*, *dcl234*, and *nrpe1-11*. *spt6lΔAh* was omitted due to low counts of $Dm^5C$. Hypermethylated sites are preferentially enriched near the 3′ end.

(F) Scatter plots correlating differential methylation ($m^5C$ difference, y-axis) with differential gene expression (DESeq2 $Log_2$ (fold change), x-axis) for the indicated mutants in comparison to WT. The $R^2$ values indicate no significant correlation between changes in $m^5C$ methylation and changes in mRNA expression.
(G) Lollipop plot of GO enrichment for genes containing hypomethylated $Dm^5C$ in *ago4-5* mutant.

(H) Metaplot profiling of DNA methylation levels (CG, CHG, CHH context) in WT plants centered on $Dm^5C$ identified as hypomethylated (left) or hypermethylated (right) in *ago4-5* mutants. The x-axis spans ±100 bp from the central $Dm^5C$. The low average methylation levels (y-axis max in CG ~5% and ~0.2% in CHH context) indicate that these specific RNA $m^5C$ sites are not strongly associated with RdDM DNA methylation.

(I, J) Motif analysis (MEME-ChIP) of sequences flanking the *ago4-5* hypermethylated (H) and hypomethylated (I) $Dm^5C$ ± 50bp. The only significantly enriched specific consensus sequence, GACUUUGW, is found for hypermethylated $Dm^5C$.

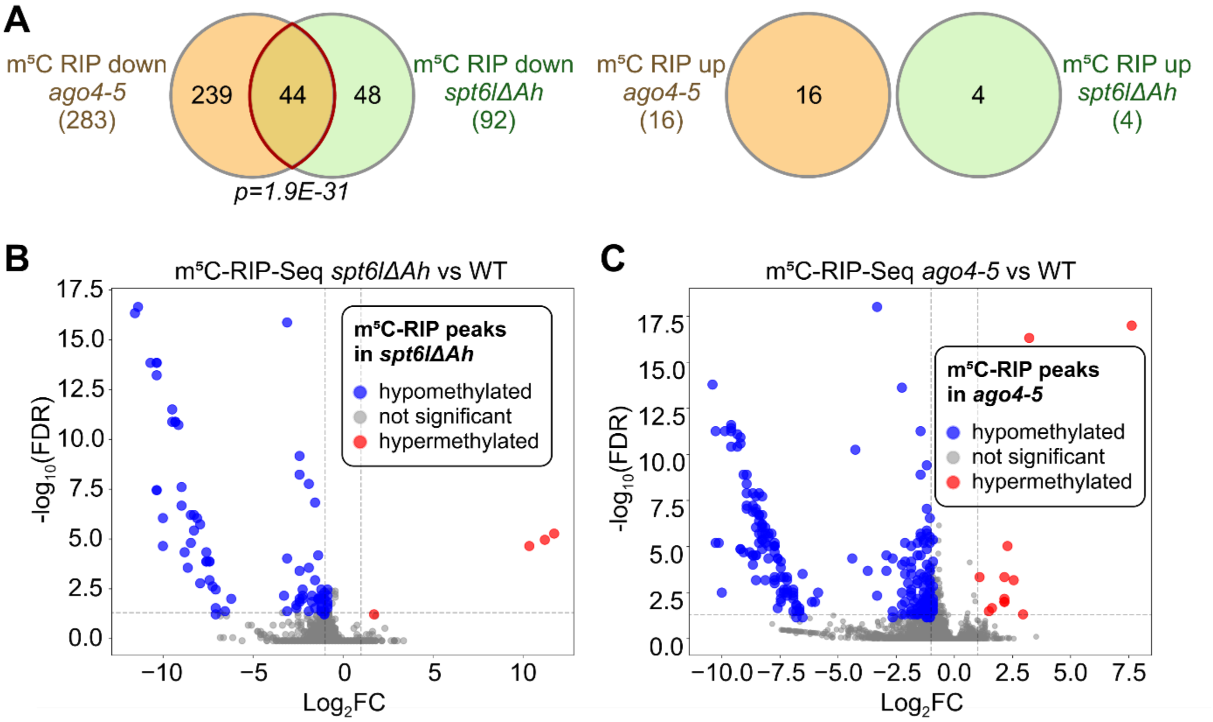


**Figure S4: Results of m$^5$C RIP-Seq confirm changes in *spt6lΔAh* and *ago4-5* mutants in RNA methylation**

(A) Venn diagrams of hypomethylated (left) and hypermethylated (right) peaks identified in *spt6lΔAh* and *ago4-5* experiments compared to WT. Statistical significance was assessed via hypergeometric probability.

(B, C) Volcano plots of differentially methylated peaks identified in the m$^5$C RIP-Seq experiment in comparison of *spt6lΔAh* and *ago4-5* compared to WT. On the x-axis is shown DESeq2 $\log_2$(fold change), and on the y-axis is significance.

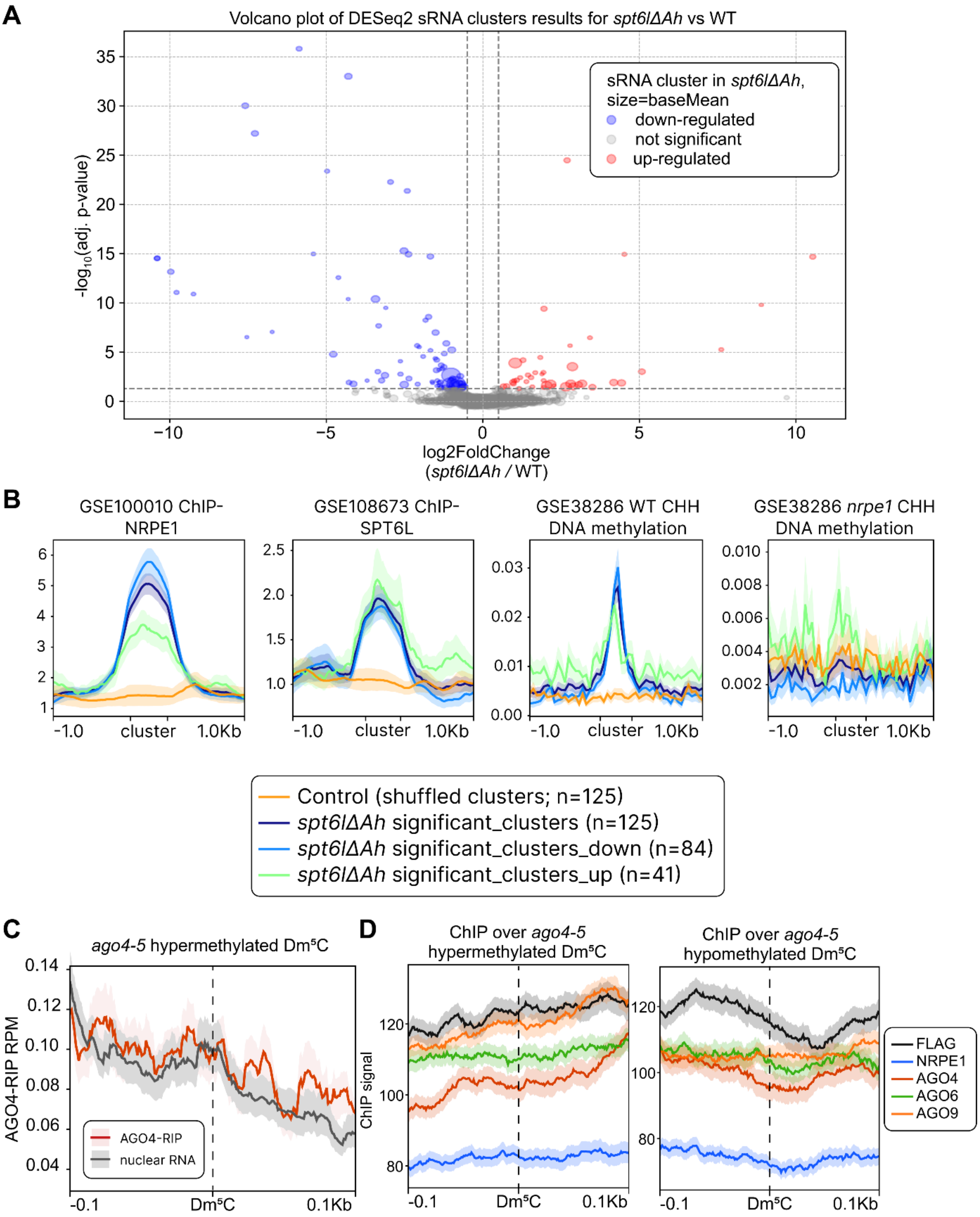


**Figure S5: SPT6L functions in RdDM independently of RNA methylation**

(A) Volcano plot displaying differential abundance of sRNA clusters in *spt6lΔAh* mutants compared to WT. On the x-axis, DESeq2 log2(fold change) is shown, and significance is shown on the y-axis. Blue dots indicate significantly downregulated clusters, red dots indicate upregulated clusters, and grey dots represent non-significant changes (adj. p-value < 0.05). The size of the dots indicates the mean read count for each cluster.

(B) Metagene profiles characterizing the genomic features of the *spt6lΔAh* differential sRNA clusters identified in (A). The panels show enrichment levels for NRPE1 ChIP-Seq (GSE100010), SPT6L ChIP-Seq (GSE108673), and CHH DNA methylation in WT and *nrpe1* backgrounds (GSE38286). The curves compare all significant clusters, downregulated clusters, upregulated clusters, and a random shuffled control set across clusters with 1.0 kb flanking regions.

(C) Metaplot analysis of AGO4-RIP sequencing data (GSE48617) compared to a nuclear total RNA as a negative control. The plot is centered on *ago4-5* hypermethylated mRNA $Dm^5C$ (n = 352) ± 100 bp. The plot depicts the mean value ± SEM.

(D) Metaplots of ChIP-seq signals for FLAG-tagged RdDM pathway components (NRPE1, AGO4, AGO6, AGO9; GSE124750) and a FLAG peptide negative control centered around *ago4-5* hypermethylated $Dm^5C$ sites (left; n = 352) and hypomethylated $Dm^5C$ sites (right; n = 257). The metaplots depict the mean value ± SEM.

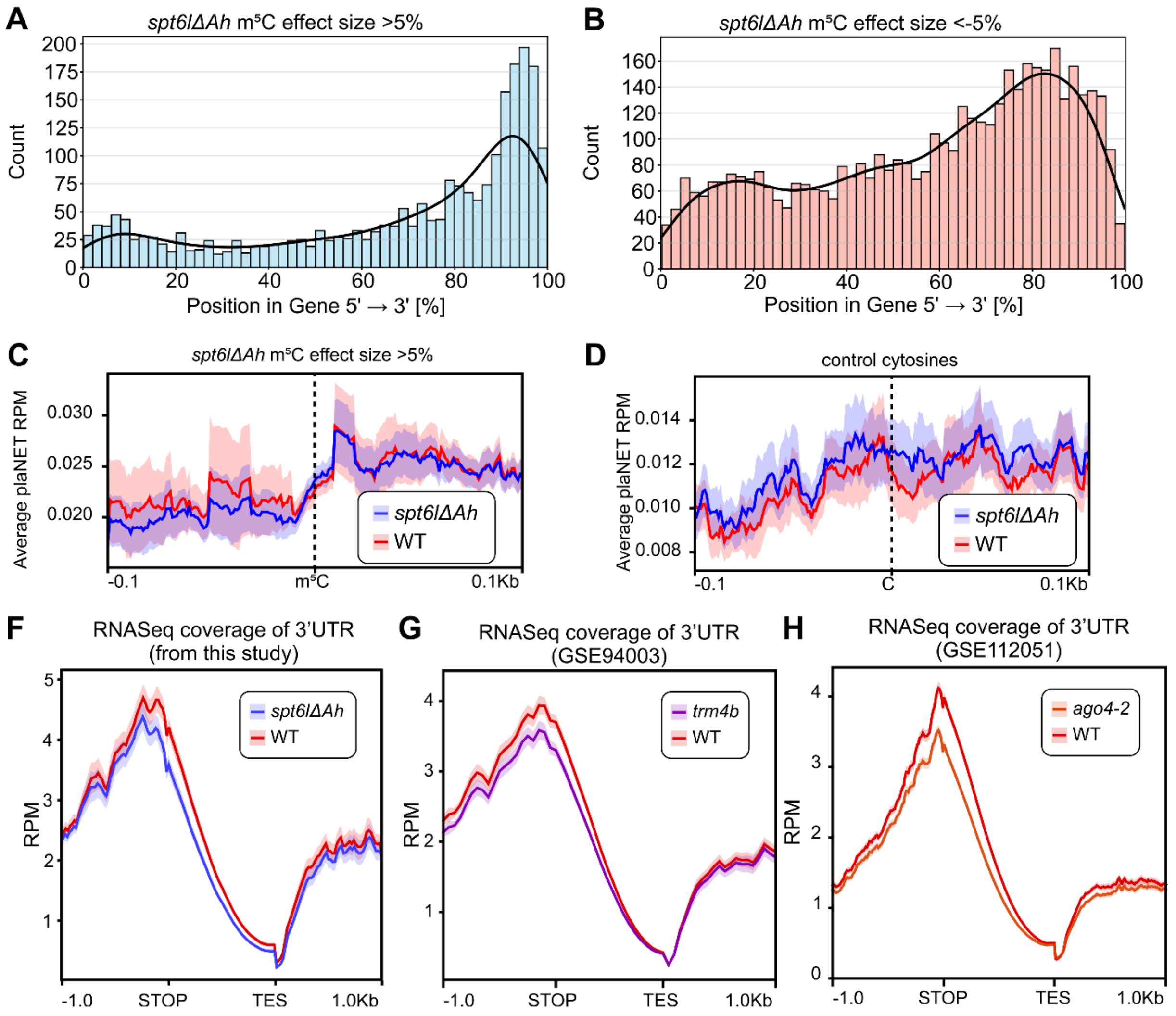


**Figure S6: The m$^5$C mark on mRNA is crucial for Pol II release for effective transcription**

(A, B) Histograms of m$^5$C positions along their corresponding genes for hypermethylated (n = 1 316; A) and hypomethylated (n = 2 400; B) m$^5$C in *spt6lΔAh*. The hypomethylated m$^5$C is mostly present in the 3' end of the genes, whereas the hypomethylated m$^5$C is distributed along the whole length of their genes.

(C, D) Metaplots of WT and *spt6lΔAh* plaNET-seq signal flanking *spt6lΔAh* hypermethylated (m$^5$C difference >5% compared to WT; n = 1 316; C), randomly selected control cytosines (n = 609; D). The lines depict mean value ± SEM as a shaded area.

(F - H) Metaplot of RNA-seq coverage (RPM) of the 3' UTR plotted from the STOP codon to the Transcription End Site (TES) for Illumina RNA-Seq for WT and *spt6lΔAh* (from this study; F), WT and *trm4b* (both from GSE94003; G), or WT and *ago4-2* (both from GSE112051; H).

**Supplementary Data S1**

Supplementary data contains results of MS/MS analysis of SPT6L(ΔAh)-FLAG samples and WT Col-0 negative control, and data of identified formaldehyde crosslinks.

**Supplementary Data S2**

Supplementary data contains results of MS/MS analysis of FLAG-AGO4 samples and WT Col-0 negative control.

**Supplementary Data S3**

DESeq2 analysis results of differentially expressed genes in *spt6lΔAh*, *ago4-5*, *dcl234*, *nrpd1*, and *nrpe1-11* compared to WT.

**Supplementary Data S4**

Supplementary data contains data of all methylatable cytosines and results of modkit analysis in *spt6lΔAh, ago4-5, dcl234, nrpd1, nrpe1-11* compared to WT, and data for the m$^5$C RIP-Seq experiment for *spt6lΔAh, ago4-5* compared to WT.

**Supplementary Data S5**

DESeq2 analysis results of differentially expressed siRNA clusters in spt6lΔAh compared to WT.

**Supplementary Data S6**

Supplementary data contain results of APAtrap analyses for *trm4b, spt6lΔAh,* and *ago4-2,* compared with their corresponding WT controls, as well as data for the Venn diagrams in Figure 5C and 5D.

**Supplementary Data S7**

Supplementary data include the oligonucleotides used in this study and the overall design of the sequencing experiments.
